# Novel Strategies for Glutamate Clearance in the Glia-Deprived Synaptic Hub of *C. elegans*

**DOI:** 10.1101/645812

**Authors:** Joyce Chan, Kirsten KyungHwa Lee, Jenny Chan Ying Wong, Paola Morocho, Itzhak Mano

## Abstract

Brain function requires the ability to form neuronal circuits that mediate focused and accurate communication. Since the vast majority of brain synapses use Glutamate (Glu) as their neurotransmitter, unintended spillover of Glu between adjacent synapses is a critical challenge. To ensure accurate neurotransmission and avert synaptic mix-up, specialized Glu Transporters (GluTs) clear the synapse of released Glu. While classical views of neuronal morphology and physiology depict isolated spiny synapses enwrapped by GluT-expressing glia, in reality, a considerable portion of synapses are flat, glial coverage in some parts of the brain is rather sparse, and extracellular space is larger than previously estimated. This suggests that diffusion in interstitial fluids might have an important role in Glu clearance in these synapses. To understand basic principles of Glu clearance in flat-, glia-deprived synapses, we study the physiology of neuronal circuits in the *C. elegans* nerve ring, the nematode’s aspiny synaptic hub. We use behavioral assays, Ca^2+^ imaging, and iGluSnFR to follow synaptic activity in intact animals. We find that synapses in a nociceptive avoidance circuit are dramatically affected by distal GluTs, while an adjacent chemoattraction circuit is controlled by proximal GluTs. We also find that pharyngeal pulsatility and mobility, which could agitate interstitial fluids, are critical for synaptic physiology. We therefore conclude that robust Glu clearance in the nematode is provided differentially by distal and proximal GluTs, aided by agitation of interstitial fluids. Such principles might be informative in determining additional factors that contribute to robust Glu clearance in other neuronal systems.

**Significance Statement:** The nervous system depends on faithful relay of information without inadvertent mixing of signals between neuronal circuits. Classical views of the nervous system depict isolated synapses, enwrapped by glia that express neurotransmitter-transporters. However, this view is incomplete, since many synapses are flat, deprived of glia, and exposed to a larger-than-expected extracellular space. We use optogenetic tools to investigate glutamate clearance strategies in the aspiny and glia-deprived synaptic hub of intact nematodes. We find a division of labor among Glutamate transporters: while some transporters display classical localization near the synapses, others are distal, and cooperate with agitation of interstitial fluids to prevent glutamate accumulation. These novel principles might contribute to synaptic clearance in higher animals, affecting normal neuronal physiology and disease.

## Introduction

Normal physiology of the nervous system depends on channeling signals into well-defined neuronal circuits, without inadvertent interruptions from neighboring circuits. This is especially difficult because 80-90% of the synapses in the mammalian brain use the same neurotransmitter, Glutamate (Glu) (Brady et al., 2012). To maintain circuit resolution and accuracy, synaptically released Glu is removed by secondary-active Glu Transporters (GluTs) (Danbolt, 2001; Tzingounis and Wadiche, 2007; Vandenberg and Ryan, 2013). While a moderate decline in GluT function blurs synaptic pulses, pronounced malfunction causes toxic accumulation of synaptic Glu, leading to neurodegeneration by excitotoxicity (seen in brain ischemia and a range of neurodegenerative diseases (Danbolt, 2001)). Robust clearance by GluTs is therefore necessary for accurate rapid signaling, curtailing spillover, and preventing excitotoxicity. Several subtypes of GluTs are found in the brain, exhibiting a range of cellular and regional expression patterns, and important differences in physiological properties. However, the functional significance of these differences remains unclear.

When considering clearance physiology, it is interesting to highlight recent insights into synaptic architecture. Classical views of synaptic organization include a presynaptic bouton, a postsynaptic spine, and GluT-expressing glia that envelope and insulate the synapse (Harris, 1999; Petralia et al., 2016). Such organization is optimal for preventing Glu spillover, while the structure of the spine is highly beneficial for creating a chemical and electrical subcellular domain (Hausser et al., 2000; Yuste, 2013). However, in critical areas of the brain, only a 1/3 of synapses are associated with glia (Ventura and Harris, 1999; Ostroff et al., 2014; Sudhof, 2018) and Glu spillover is prominent (Kullmann and Asztely, 1998). Furthermore, some synapses, especially in the developing brain, are flat shaft synapses (Segal, 2010) that have distinctive functional features, allowing for more direct passive cable conductance of depolarization (Rall, 1959; Magee, 2000). Flat shaft synapses show scant GluTs (Chaudhry et al., 1995), relying more on diffusion as the primary mode of Glu clearance (Barbour et al., 1994; Clements, 1996; Thomas et al., 2011). Furthermore, recent studies suggest that the fraction of extracellular space in the intact brain is much higher than previously suggested, particularly around synaptic connections (Korogod et al., 2015; Tonnesen et al., 2018). Additional studies emphasize the significance of the brain’s glymphatic system and its ability to clear macromolecules by bulk flow (Tarasoff-Conway et al., 2015; Da Mesquita et al., 2018; Rasmussen et al., 2018). Functional models suggest that an open synaptic architecture is conducive to clearance by diffusion and bulk flow (Nicholson and Hrabetova, 2017). Nonetheless, clearance of Glu in aspiny, glia-deprived synaptic hubs exposed to high-fraction extracellular space and interstitial fluids remains understudied. How circuit coherence is maintained in the absence of anatomical synapse isolation remains poorly understood.

To address this gap in our knowledge, we turn to the model system of the transparent nematode *C. elegans*, which combines powerful genetic, molecular, and intact-animal optogenetic tools, with an extraordinarily detailed description of the nervous system (White et al., 1986). In the nematode nerve ring (its hub of synaptic activity), aspiny neurons encircle the animal’s pharynx and form multiple *en passant* synapses between neurites. Communication relies mostly on passive cable propagation (Goodman et al., 1998). Four cephalic sheath glia cells wrap around the circumference of the nerve ring (Altun et al., 2002-2019) and (for the most part, with some exceptions (Singhvi and Shaham, 2019)) do not provide separation between synapses, or separate interstitial fluids in the neuropil from the body fluids of the pseudocoelon. Since Glu is a key excitatory neurotransmitter in *C. elegans* (Brockie and Maricq, 2006), the nematode maintains a fully functional Glu signaling system and high fidelity circuit resolution under an unfavorable nervous system configuration, one that lacks anatomical separation and utilizes the same neurotransmitter in many adjacent synapses. In our studies we ask if these challenges are addressed in *C. elegans* by a compensatory, especially robust function of the Glu clearance system to maintain the accuracy of Glu signaling and circuit resolution.

## Materials and methods

### Strains and maintenance

All *Caenorhabditis elegans* strains were cultured at 20°C on MYOB plates (Brenner, 1974; Church et al., 1995) with *Escherichia coli* strain OP50 as food source. Our wild type strain is Bristol N2. Other strains used for this study are: For behavioral analysis: VM1268: *nmr-1(ak4) II; glr-2(ak10), glr-1(ky176) III*; ZB1113: *glt-1(ok206) X*; ZB1096: *glt-3(bz34) IV*; ZB1098: *glt-4(bz69) X*; IMN16: *glt-3(bz34), glt-6(tm1316), glt-7(tm1641) IV*; For GCaMP imaging in AVA: QW625 (*lin-15; zfIs42*[*P_rig-3_::GCaMP3::SL2::mCherry; lin-15(+)*]); IMN18: *glt-1(ok206) X; zfIs42*; IMN19: *glt-3(bz34), glt-6(tm1316), glt-7(tm1641) IV; zfIs42*; IMN20: *glt-4(bz69) X, zfIs42*; For GCaMP imaging in ASH: CX10979 (*kyEx2865*[*P_sra-6_::GCaMP3; ofm-1::gfp*]); For iGluSnFR imaging around AVA’s nerve ring neurites: CX14652: *kyEx4787* [*P_rig-3_::iGluSnFR, unc-122::dsRed*]; IMN50: *glt-1(ok206) X; kyEx4787*; IMN51: *glt-3(bz34), glt-6 (tm1316), glt-7 (tm1641) IV; kyEx4787*. Details of the *glt* mutant strains were previously described (Mano et al., 2007). The strains carrying *glt* mutations and AVA neurons expressing GCaMP or iGluSnFR (IMN18, IMN19, IMN20, IMN50 and IMN51) were generated by crosses between the corresponding *glt* mutants and the QW625 or CX14652 strain. QW625 (Shipley et al., 2014) was a gift from the Alkema lab (U. Mass. Med. Sch.) and was obtained via the Biron lab (U. Chicago) (Iwanir et al., 2013). The CX10979 (Larsch et al., 2013) and CX14652 (Marvin et al., 2013) strains were gifts from the Bargmann lab (Rockefeller U.).

### Behavioral assays

Spontaneous mobility assay (Zheng et al., 1999; Brockie et al., 2001a; Mellem et al., 2002) and nose touch assay (Kaplan and Horvitz, 1993; Hart et al., 1995) were conducted as previously described. For the spontaneous mobility assay, the forward locomotion duration was determined until the individual worm either halts movement or reverses. For nose touch response we determined the number of worms responding (halting or reversing) or not responding (continuing forward mobility) to collision with an eyelash. For each session, about 30 animals of each genotype were tested.

Avoidance drop assay was conducted for testing worms’ avoidance response to a range of NaCl concentrations as previously described (Chatzigeorgiou et al., 2013). We tested NaCl concentrations, ranging from 0.1 mM to 1000 mM, dissolved in 1mM MgSO_4_, 1 mM CaCl_2_, and 5 mM KPO_4_. For each session, worms were transferred to an unseeded MYOB plate and left there for 15 min to remove remaining food on their body. Worms were then transferred to the assay plate and were given an additional 15 min to adjust to the buffer solution. A capillary tube was used to make a drop near the tail of a forward-moving worm. The drop immediately surrounds the whole worm body. A mere mechanosensory stimulation to the tail will stimulate the worm to rush forward, but chemorepellent properties of solutes sensed in the nose will cause the worm to halt or reverse. The worms’ response within 4 s after the drop was denoted as 0 if the worms continued their forward movement, and as 1 if worms either stopped or reversed. To present the trend of avoidance over a range of NaCl concentrations, we used avoidance index equal to the percentage of worms avoiding specific NaCl concentration.

### Microfluidics and imaging

Prior to imaging, adult worms were first transferred onto non-seeded MYOB holding plate. Worms were allowed to traverse the holding plate for 10 minutes, removing excess bacteria adhering to their bodies. For stimulation experiments with glycerol, S-basal buffer was added onto the holding plates for no more than 10 minutes before positioning of each individual worm into the worm channel of the microfluidics chip. For the salt stimulation experiments, a solution consisting of 1 mM MgSO_4_, 1 mM CaCl_2_, and 5 mM KPO_4_ (salt stimulation buffer) was used instead (Oda et al., 2011).

The Chronis & Bargmann worm behavioral chip (Purchased from MicroKosmos) was used for GCaMP and iGluSnFR imaging experiments (Chronis et al., 2007). This system allows for temporal stimulation of amphid sensory neurons located on the worm nose. Worms were physically restrained in a specially fitted channel (the worm trap) designed to hold adults without the use of paralytic agents that may interfere with normal physiology. A system of liquid streams controlled under laminar flow were manipulated to present either control buffer or stimulus to the worm nose (Chronis et al., 2007) via a three-way electric valve (Lee Company). GCaMP and iGluSnFR transients from live nematodes trapped in the chip were recorded as previously described by Chronis & Bargmann. To generate the stimulant solutions, glycerol was dissolved in S Basal buffer to a final concentration of 1 M (Chronis et al., 2007), while 1 mM NaCl was dissolved in salt stimulation buffer. For paralysis experiments, either 2 mM tetramisole (Sigma) or 0.3 M BDM (2,3-Butanedione monoxime) (VWR) dissolved in either S-basal or salt stimulation buffer was introduced via the buffer channel of the microfluidics system to paralyze the anterior portion of the worm. Paralysis was induced by exposing the head of the immobilized animals contained in the worm trap of the microfluidics chamber to the paralytic agents for no more than 10 minutes.

Our imaging system consists of a Zeiss Axiovert 200 M motorized inverted microscope, Lumencor SOLA solid state white light source, Ludl filter wheel controller, Q Imaging EXiTM Blue camera, and ValveBank4 controller (AutoMate). Metamorph software (Molecular Devices) was used for image processing and acquisition. GCaMP transients were captured with a 63x objective lens (10 frames/s), while iGluSnFR transients were captured at 40x magnification (3 frames/s) for each experiment duration.

We used ΔF/F to indicate change in fluorescence intensity. F was defined as the baseline fluorescence intensity of AVA during a period of either 3 seconds (for recording effects of salt stimulation in AVA GCaMP and the iGluSnFR experiments) or 4 seconds (for recording AVA GCaMP signals during response to glycerol). Intensity measurements were restricted to AVA cell body for GCaMP imaging, and neuronal processes for iGluSnFR imaging. This was achieved by first setting an inclusive intensity threshold to define the range of fluorescence to capture (i.e., neuronal soma or process, which has higher intensity compared to other cells and structures), then defining a region of interest (ROI) to capture from. To restrict iGluSnFR measurements to AVA neuronal processes in the absence of animal paralysis, the ROI was manually shifted to follow the location of the neurite, and fluorescence was logged manually. Reporter transients were analyzed by comparing subsequent intensity readouts to the baseline intensity, expressed as a change in percentage.

### Statistical analysis

All statistical analyses was performed utilizing GraphPad Prism software. For the nose touch and the drop test assays, we used the Student’s t-test with Welch’s correction to compare significance of differences between mutant and N2 control strains. For GCaMP and iGluSnFR data, ANOVA with post hoc Bonferroni test was used for multiple group comparison of their means. Error bars denote SEM, and statistical significance is noted with * based on P values.

## Results

### Behaviors mediated by the avoidance circuit are strongly sensitive to manipulation of distal, but not proximal, GluTs

Our previous studies of Glu clearance in *C. elegans* suggested an unusual functional organization of Glu uptake, emphasizing the role of distal clearance (Mano et al., 2007). GFP-promoter fusion experiments reveal the expression of two GluT genes close to the nerve ring: *glt-1* is expressed in head muscles and in the hypodermis (Katz & Shaham later show glial expression of *glt-1* (Katz et al., 2018)), while *glt-4* is expressed in some head neurons. The three remaining highly expressed GluT genes (*glt-3, glt-6*, & *glt-7*) are expressed distally from the synapses, solely on the canal cell (an H-shaped tubular cell running along the length of the animal). This structure passes the nerve ring latero-ventrally, maintaining a distance of ~10 microns from most glutamatergic synapses. We therefore classified *glt-1* & *glt-4* as proximal GluTs, and *glt-3, glt-6,* & *glt-7* as distal GluTs. Amino acid sequences at the GluT active site (conserved across phyla and in nematode proximal GluTs) suggest that the nematodes’ distal GluTs may have unusual physiological properties. Despite these unusual characteristics of distal GluTs, our genetic studies have shown that these GluTs have a critical effect on Glu synapses under both normal (Mano et al., 2007) and pathological conditions (Mano and Driscoll, 2009; Mojsilovic-Petrovic et al., 2009; Tehrani et al., 2014; Del Rosario et al., 2015).

We now examine the functional role of these distal GluTs in more detail. We produced a combined knockout of all distal GluTs (*glt-3, glt-6,* & *glt-7*) but were unable to produce a combined KO of proximal GluTs (*glt-1*; *glt-4*), potentially because of a vital role in early development. We find that KO of all distal (but not proximal) GluTs shortens the duration of spontaneous forward mobility and disrupts nose touch response (Figure 1). Both behaviors are known to be mediated by a bilateral pair of glutamatergic polymodal nociceptive sensory neurons ASH (L/R), and their main GluR-expressing postsynaptic target neurons AVA, AVD, & AVE (which mediate animal reversal, and are therefore denoted as the nociceptive or avoidance circuit; minor additional/indirect contributions come from additional neurons) (Mellem et al., 2002; Brockie and Maricq, 2006). The balance between forward and backward runs in spontaneous mobility (quantified as either the duration of forward runs before halting, or the frequency of reversals) and sensitivity to nose touch (quantified as the fraction of animals responding to a collision with an obstacle) are considered to be very sensitive measures of glutamatergic activity in these synapses (Zheng et al., 1999; Burbea et al., 2002; Chang and Rongo, 2005; Brockie and Maricq, 2006). The behavioral effect of combined distal GluT KO is in line with our previous analyses of *glt-3* (a single KO strain), and the ability of that mutation to compensate for other mutations (such as *eat-4* and *glr-1*) that reduce Glu signaling in the avoidance circuit (Mano et al., 2007). Therefore, our observations on behaviors of distal GluT mutants (but not proximal GluT) mutants in response to spontaneous mobility and nose touch are consistent with a model where *glt-3, glt-6*, & *glt-7* KO caused an accumulation of Glu in the ASH -> AVA/AVD/AVE synapses.

**Figure 1:**
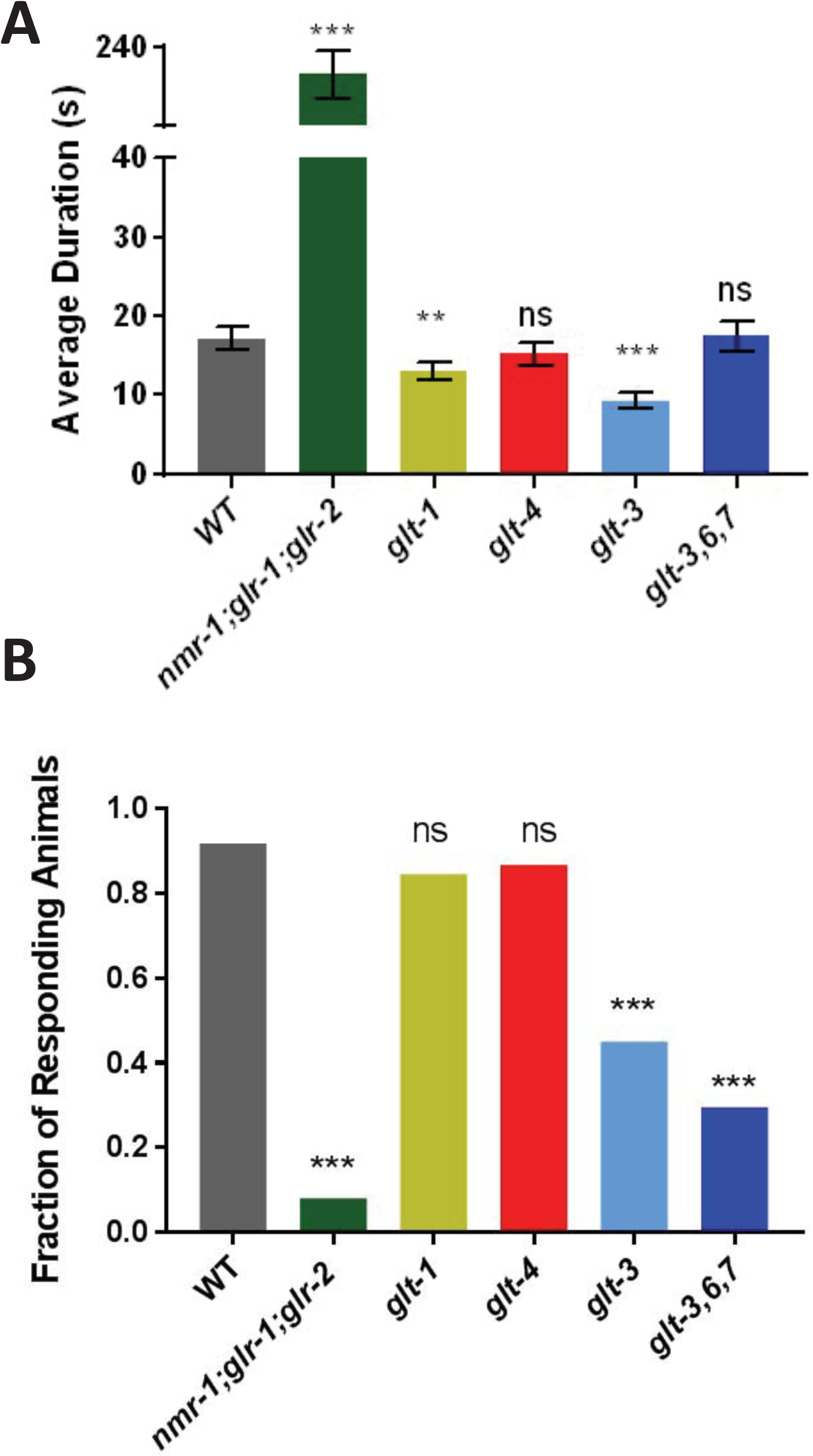
Distal GluT KOs disrupt behaviors mediated by ASH -> AVA avoidance circuit. (A) Duration of forward runs in spontaneous mobility assay. *glt-3* (distal GluT KO) animals are impaired in forward run duration, although the duration of forward run of proximal GluT KO animals (*glt-1,*or *glt-4*) is intact. The *nmr-1; glr-2;glr-1* strain (where all ionotropic GluR responses of the avoidance circuit are eliminated) is used as a control where Glu signaling is reduced, resulting in excessive forward mobility. (B) Nose touch response of distal GluT KOs, *glt-3* and *glt-3, glt-6, glt-7* KO is reduced. Colored bars represent the fraction of animals responding to nose touch. Both underactive (*nmr-1; glr-2; glr-1* animals) and overactive (*glt-3 and glt-3, glt-6, glt-7* animals) Glu signaling causes impaired response to nose touch. Significance of differences from control (WT, N2 strain) mean (A) or distribution (B) is indicated by asterisks. ** *P* = 0.10; *** *P* < 0.001. Student’s t-test with Welch’s correction was used in both (A) and (B). Error bars indicate SEM; n = 15-120 per genotype (A), n = 45-90 per genotype (B).

### GCaMP- and iGluSnFR-based imaging of neuronal activity in AVA provide direct evidence for hyper stimulation of the avoidance circuit in distal GluT KO animals

To examine the effect of GluT KO on physiological responses of the postsynaptic neurons in the avoidance circuit more directly, we set up a nematode microfluidics and imaging system similar to the setup pioneered by the Bargmann lab (Chalasani et al., 2007; Chronis et al., 2007; Chalasani et al., 2010; Chronis, 2010). In this setup, the worm is restrained in a microfluidic chip in absence of paralytic agents, restricting exposure of stimulant solutions to the nose (and consequently, sensory amphid neuron endings). The nematode’s transparency allows for non-invasive, *in vivo* recording of physiological neuronal activity, which we monitor using genetically encoded fluorescent Ca^2+^ sensor (GCaMP) expressed in identified neurons. In summary, this system provides us an easily accessible window into neuronal activity in specific postsynaptic neurons in intact animals without compromising the hydrodynamics of interstitial fluids or changes in concentrations of our neurotransmitter of interest. We used GCaMP expressed in AVA (the main mediator of avoidance responses) to monitor the activity of this neuron in response to ASH stimulation by high concentration glycerol (Figure 2, measuring changes in whole-soma GCaMP fluorescence, which is correlated with membrane depolarization in this cell (Piggott et al., 2011)). We used the QW625 (*zfIs42*[*P_rig-3_::GCaMP3::SL2::mCherry*]) strain (Shipley et al., 2014) due to its inter-animal consistency (as a strain containing a genetically integrated reporter) in GCaMP intensity. Though ASH-specific nociceptive stimuli are known to produce measurable GCaMP responses in AVA in freely moving animals (Chronis et al., 2007; Piggott et al., 2011; Cho and Sternberg, 2014), the consistency of this response is reduced in animals restrained in the microfluidic chip (in line with observations made by other labs on AVA responses in restrained vs free animals, S. Chalasani, personal communication). A small AVA response is observed when the proximal GluTs *glt-1* or *glt-4* are eliminated. In sharp contrast, elimination of the distal GluTs *glt-3, glt-6*, & *glt-7* elicited a very prominent response to glycerol (Figure 2).

**Figure 2:**
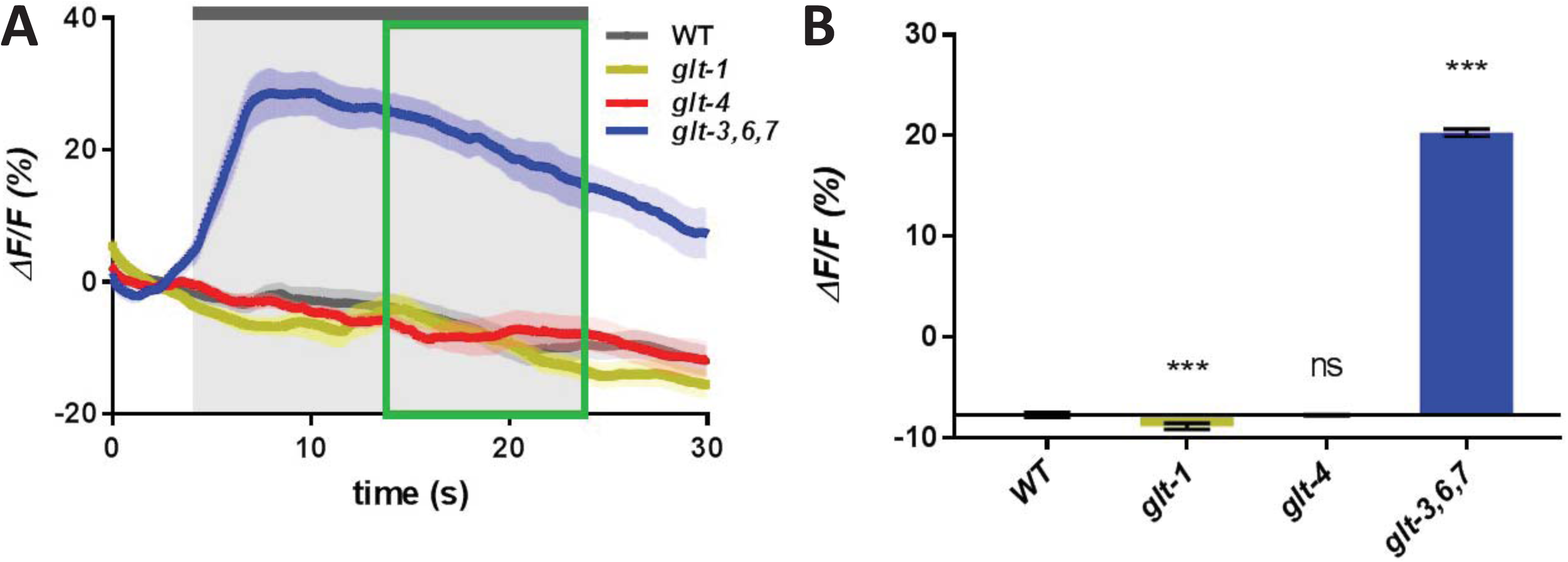
Distal GluT KOs enhance Glu signaling in ASH -> AVA synapse. Glycerol-induced calcium responses in AVA neurons of WT and GluT KO animals. **(A)** Average traces of changes in GCaMP3 fluorescence. Shaded areas above and below the trace represent SEM. The fluorescence intensity of the first 4 s of the recording was averaged to serve as the baseline fluorescence, F_0_. Beginning at *t* = 4 s, light gray shading indicates 20 s period of exposure to a 1 M glycerol stimulus. Within this exposure period, green box indicates a window of 10 s, after neural response to stimulation reached a relative stability, for which the steady-state fluorescence change (ΔF) was averaged and presented in the bar graphs. **(B)** Average steady-state response of each strain is compared to the average steady state response of WT animals (indicated as a horizontal line). Error bars represent SEM. *** *P* < 0.001, ANOVA with Bonferroni correction. n = 15 for each strain.

Next, we sought to determine if the increased responses in AVA in distal GluT KO animals arose directly from increased concentration of perisynaptic Glu around AVA’s neurites, and not from heightened response sensitivity of AVA, or from gap junctions from other neurons affecting AVA. To directly track effects on perisynaptic Glu concentrations we used the Glu-sensitive extracellular sensor iGluSnFR, expressed specifically in AVA neurons in CX14652 (*kyEx4787* [*P_rig-3_::iGluSnFR, unc-122::dsRed*]) animals (Marvin et al., 2013), and measured changes in florescence on AVA’s ring neurites. Measuring changes in iGluSnFR fluorescence in AVA neurites in the nerve ring is technically challenging, especially when the animal continues to perform pharyngeal pumping and local pharynx movements in the absence of pharmacological paralyzing agents. Nonetheless, in response to ASH stimulation we saw a small increase in Glu readouts in AVA’s nerve ring neurites in *glt-1* mutant and a dramatic increase in *glt-3* mutants (Figure 3).

**Figure 3:**
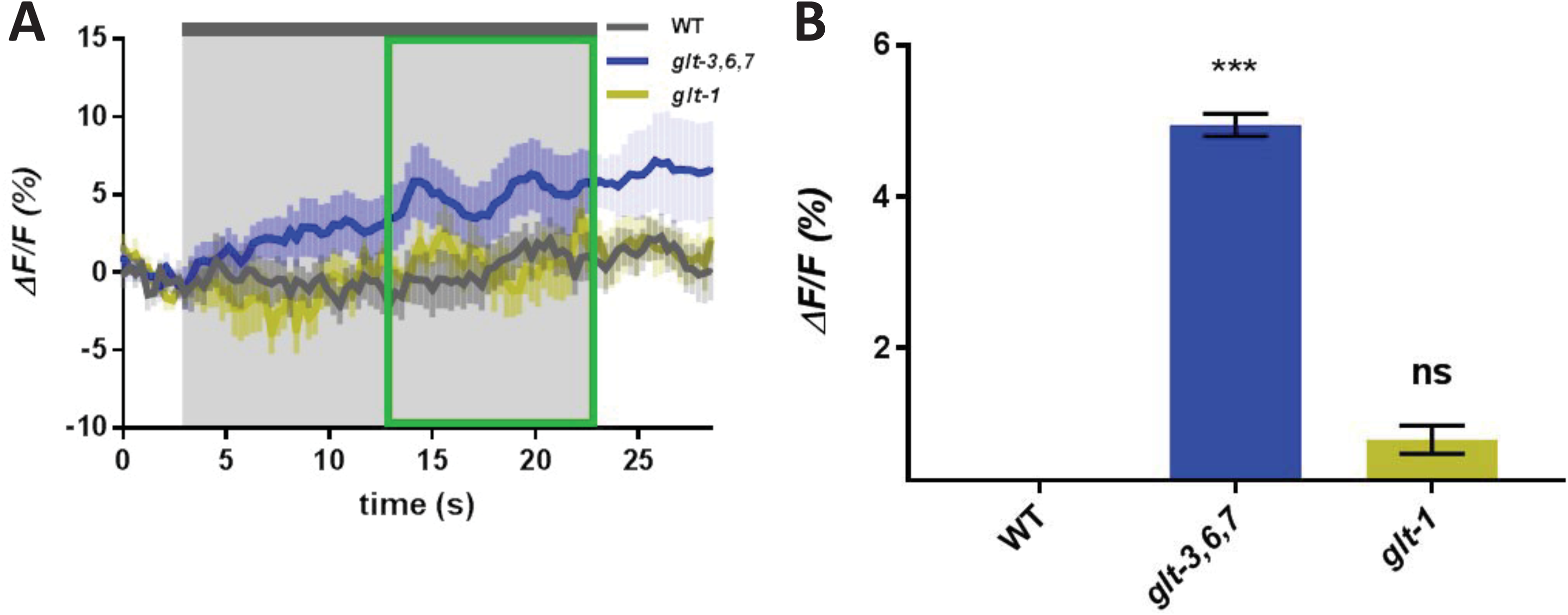
Distal GluT KOs promote accumulation of Glu at the AVA synapse following 1M glycerol stimulation. Glycerol-induced changes in Glu concentrations in nerve ring processes of AVA neurons in WT and GluT KO animals. Average traces of changes in iGluSnFR fluorescence and averaged steady-state responses to a 1 M glycerol stimulus are analyzed similarly to those in Figure 2. Error bars represent SEM. \*\*\**P* < 0.001, ANOVA with Bonferroni correction.. n = 5-6 for each strain.

The GCaMP and iGluSnFR–based imaging data correlate with the behavioral observations, and provides direct evidence (at the level of perisynaptic Glu concentrations and postsynaptic response physiology) that elimination of distal Glu clearance strongly potentiates Glu signaling between ASH and AVA in the avoidance circuit, while elimination of proximal GluTs produces only a small potentiation of AVA responses. Altogether, these observations support that, unlike proximal GluTs, distal GluTs have a major role in clearing Glu from ASH -> AVA synapses. Given the challenges in measuring changes in perisynaptic Glu concentrations with iGluSnFR in non-paralyzed animals, we proceeded to further analysis mostly by using GCaMP imaging to detect changes in intracellular Ca^2+^ concentrations in the soma as a reasonable reporter of synaptic excitation. As we start gaining insight into the preferential contribution of distal vs proximal GluT to Glu clearance in the avoidance circuit, we became interested to compare it to the contribution of different GluTs to the activity of other circuits, such as a circuit that normally mediates chemoattraction.

### Reduction of proximal Glu clearance generates avoidance response to low concentration of NaCl

We next assessed the effect of GluT KO on the nematode’s response to NaCl, since NaCl affects a pair of functionally well-separated but anatomically adjacent neuronal circuits. The response to NaCl is mediated in the nematode mostly by two pairs of sensory neurons, depending on concentration (Bargmann and Horvitz, 1991; Hukema et al., 2006; Suzuki et al., 2008; Oda et al., 2011; Chatzigeorgiou et al., 2013; Kunitomo et al., 2013; Leinwand and Chalasani, 2013). Low NaCl concentrations (< 200 mM) are chemoattractive: the bilateral ASE sensory neurons are sensitive to low NaCl concentrations and use Glu to signal forward mobility by regulating AIB, AIA, and AIY interneurons. Combined with alternating head movements, ASEL detects an increase in salt concentration, inhibiting AIBs through Glu-gated Cl^-^ channels such as GLC-3 to suppress reversal. ASER detects a concentration decrease, stimulating the AIBs through AMPA-Rs (such as GLR-1) and metabotropic GluRs, causing animal reversal (Suzuki et al., 2008; Wang et al., 2017; Kuramochi and Doi, 2018). In contrast to the sensitivity of ASEs to low salt concentrations, high NaCl concentrations are nociceptive and generate avoidance responses, stimulating the ASH neurons and their associated avoidance circuit (Chatzigeorgiou et al., 2013). Since the synapses of avoidance and chemoattractive circuits are localized with considerable anatomical proximity in the nerve ring (see below), we sought to determine whether GluTs contribute to the maintenance of these circuits’ functional separation, and if GluT KO might cause Glu spillover between these circuits. As a first attempt to find evidence that such a spillover occurs, we tested the possibility that low NaCl concentrations (which normally stimulate the ASE->AIA/AIB/AIY chemoattractive circuit) might trans-activate the ASH->AVA/AVD/AVE circuit and produce avoidance when GluTs are absent. We therefore measured the concentration dependence of avoidance response to NaCl using the drop assay (Suzuki et al., 2008; Chatzigeorgiou et al., 2013). We found that mutations in *glt-1* or *glt-4* (the two proximal GluTs), but not *glt-3, glt-6*, & *glt-7* KO (the distal GluTs) dramatically changed the normal behavioral dose-response curve of NaCl: an extremely low NaCl concentration (1mM), which is normally attractive, is chemorepulsive in proximal GluT mutants (Figure 4). This result suggests that Glu from the ASE -> AIA/AIB/AIY circuit might have spilled over to the ASH -> AVA/AVD/AVE circuit.

**Figure 4:**
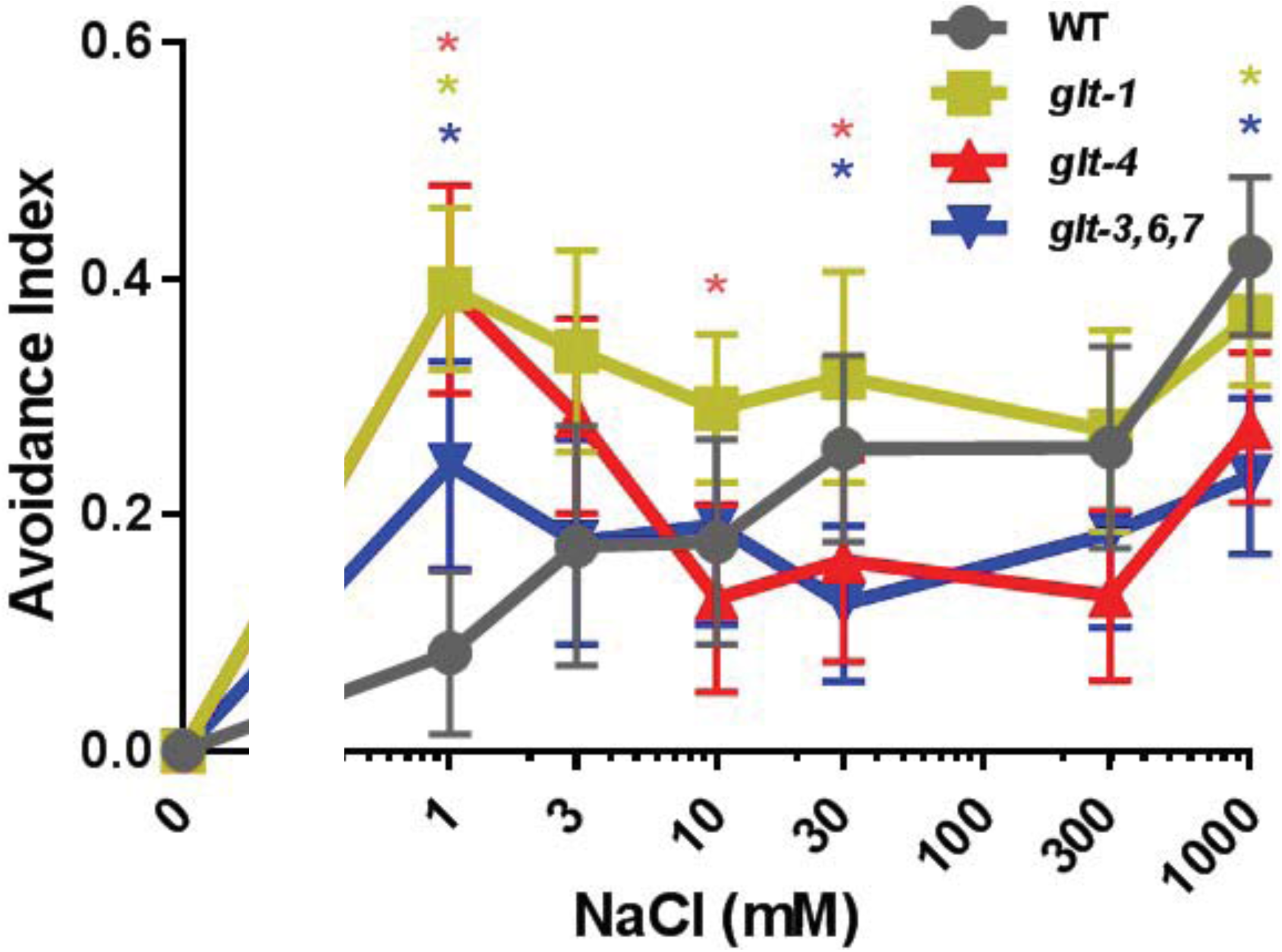
Absence of proximal GluTs abnormally switches worms response to low concentration NaCl from attraction to avoidance. A drop of buffer containing the indicated concentration of NaCl was presented to the tail of a forward-moving worm. Animals presented either an avoidance response (stopping/reversing) or an attraction response (continuing forward traversal). Avoidance index is the percentage of worms that presented an avoidance response. Difference from control (WT) distribution at each NaCl concentration is indicated by asterisks. * *P* < 0.05, Chi-square test. Error bars indicate SEM; n = 11-62 for each data point.

While spillover of Glu between these two circuits is one possible explanation for our observations, a battery of other explanations could be offered. One alternative explanation is that indirect signaling between the two circuits through other neuronal connections. However, the contribution of indirect inputs to AVA from neurons such as AIB is considered relatively minor (see WormWeb.org). Another possibility is that the ASH neurons are in fact sensitive to low NaCl concentration and release weak Glu signals onto AVA that can be potentiated by GluT KO. To negate the latter possibility, we directly monitored the activation of ASH by measuring its Ca^2+^ responses to different concentrations of NaCl, using the CX6632 *kyEX728* [*P_sra-6_::GCaMP*] strain (Chronis et al., 2007). In accordance with other reports (Leinwand and Chalasani, 2013), we find that ASH responds only to very high NaCl concentrations (Supplementary Figure 1). Altogether, these observations further suggest that the avoidance response to 1mM NaCl in proximal GluT KOs is not due to potentiation of previously weak presynaptic release from ASH neurons (that, until recently, went undetected by postsynaptic measurement), and is therefore more likely to represent Glu spillover from other sensory neurons.

### Reduction of proximal Glu clearance causes abnormal AVA activation in response to low NaCl concentration

The glutamatergic sensory ASE neuron was previously shown to be responsible for responses to low NaCl concentrations by triggering activity in the ASE -> AIA/AIB/AIY chemoattractive circuit. The ASE -> AIA/AIB/AIY circuit is therefore the leading candidate to be the source of Glu in the abnormal avoidance responses we observe in proximal GluT KO animals under stimulation with low NaCl concentrations. We then progressed from the behavioral to the physiological level, recording Ca^2+^ responses to low NaCl concentrations in AVA neurons in GluT KO animals. We find that AVA’s neural activity correlates with the avoidance responses in the NaCl drop assay (Figure 5): while low NaCl concentrations do not elicit an AVA response in either WT or distal GluT (*glt-3, glt-6*, & *glt-7*) KO background, reducing proximal Glu clearance by either *glt-1* KO or *glt-4* KO results in a robust AVA Ca^2+^ response to stimulation of animals by low NaCl. Again, we verified that the somatic Ca^2+^ responses from GCaMP recordings are correlated with perisynaptic increases in Glu levels, as recorded with iGluSnFR in AVA neurites (Supplementary Figure 2). These observations suggest that in the absence of proximal GluTs, Glu released by circuits sensitive to low NaCl concentrations now reach AVA. The most direct explanation that suffices to account for these observations is that in the absence of proximal GluTs, Glu released from ASE (in response to animal stimulation with low NaCl concentration) escapes the ASE -> AIA/AIB/AIY circuit and spills over to the ASH -> AVA/AVD/AVE circuit (though we do not exclude the alternative possibility of potentiation of inputs to AVA from other, indirect connections). According to this view, Glu spilled over from ASE-AIA/AIB/AIY synapses (or, in the alternative model, Glu coming from irregularly potentiated indirect connections) now generates an abnormal avoidance response to a normally chemoattractive stimulant. These observations also suggest that in the WT, proximal GluTs are positioned to have a privileged effect on Glu released by ASE and prevent its spillover beyond its designated postsynaptic targets, while distal GluT preferentially clears Glu released from ASH. Therefore, different GluTs exhibit specialized or privileged functional roles in different synapses.

**Figure 5:**
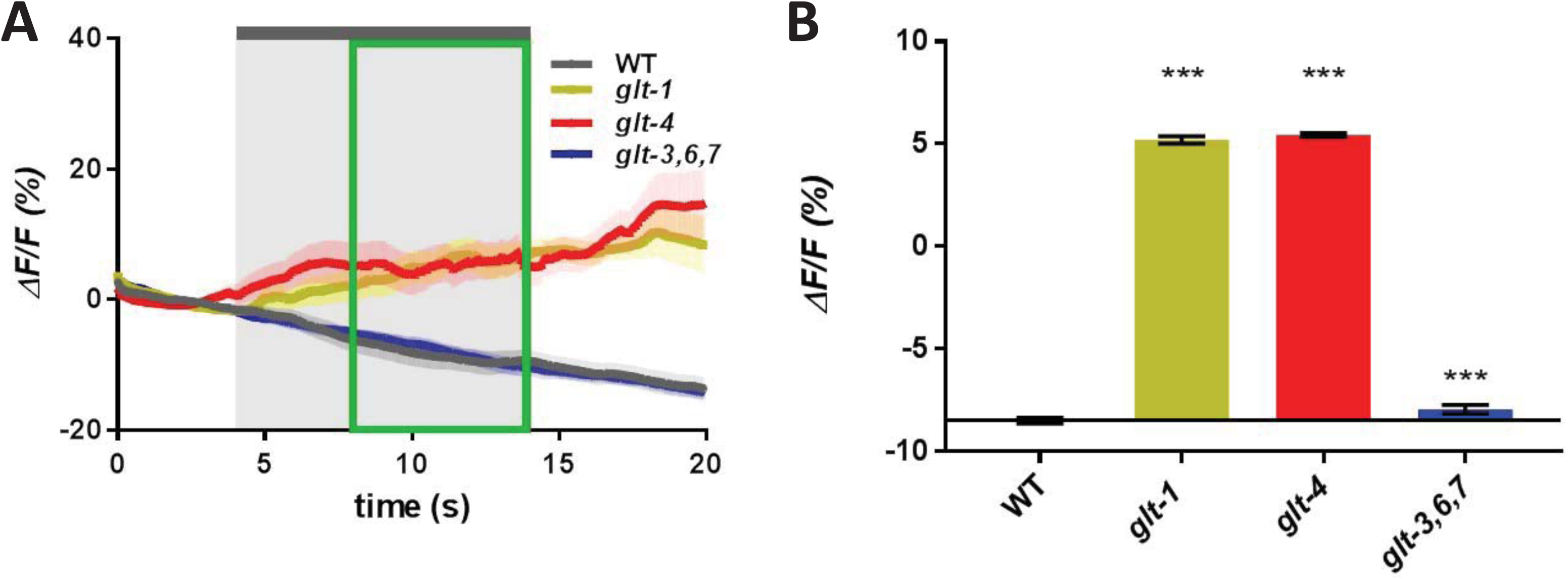
Proximal GluT KOs cause AVA to respond to low NaCl concentrations. Low-salt-induced calcium responses in AVA neurons of WT and GluT KO animals. Average traces of changes in GCaMP3 fluorescence and averaged steady-state responses to a 1 mM NaCl stimulus are analyzed similarly to those in Figure 2. *** *P* < 0.001, ANOVA with Bonferroni correction. n = 15-16 for each strain.

### Differential localization of ASH and ASE synapses in the nerve ring might contribute to the differential effect of proximal and distal Glu clearance

Intrigued by the privileged role of different GluTs in different circuits, we looked into existing anatomical data on the location of synapses in the ASE -> AIA/AIB/AIY chemoattraction circuit and the ASH -> AVA/AVD/AVE avoidance circuit. The nematode connectome was established by utilizing EM images of coronal sections (presented as consecutively-numbered slices) and reconstruction data (initially published as “The Mind of the Worm”) (White et al., 1986). This data was later developed by the Hall and Emmons labs in the form of the comprehensive web resources available at WormAtlas.org, WormWiring.org, and CytoShow.org (Altun et al., 2002-2019; Jarrell et al., 2012). The data is now available as a collection of fully annotated consecutive slices and as a complete record of the location and identity of the synapses, as well as 3D reconstruction of all neurons (Brittin et al., 2018). We used these web resources to locate the synapses relevant to our study. We find that the neuronal processes of the two circuits we study are indeed adjacent, as they track rather closely in the nerve ring. Furthermore, the ASE -> AIA/AIB/AIY and ASH -> AVA/AVD/AVE synapses, as identified in the Worm Connectome project, are frequently found in close proximity to each other. Figure 6 shows a detail from slice #87 of the EM series, where ASE -> AIB/AIY synapses (cell processes labeled with red outline) and ASH -> AVD synapses (cell processes labeled with blue outline) are found in close proximity, with the synapse of the latter gradually expanding to add AVA to the left of AVD (fully visible in slice #91). Overall, the synapses of the two circuits are positioned favorably enough for Glu spillover to occur.

**Figure 6:**
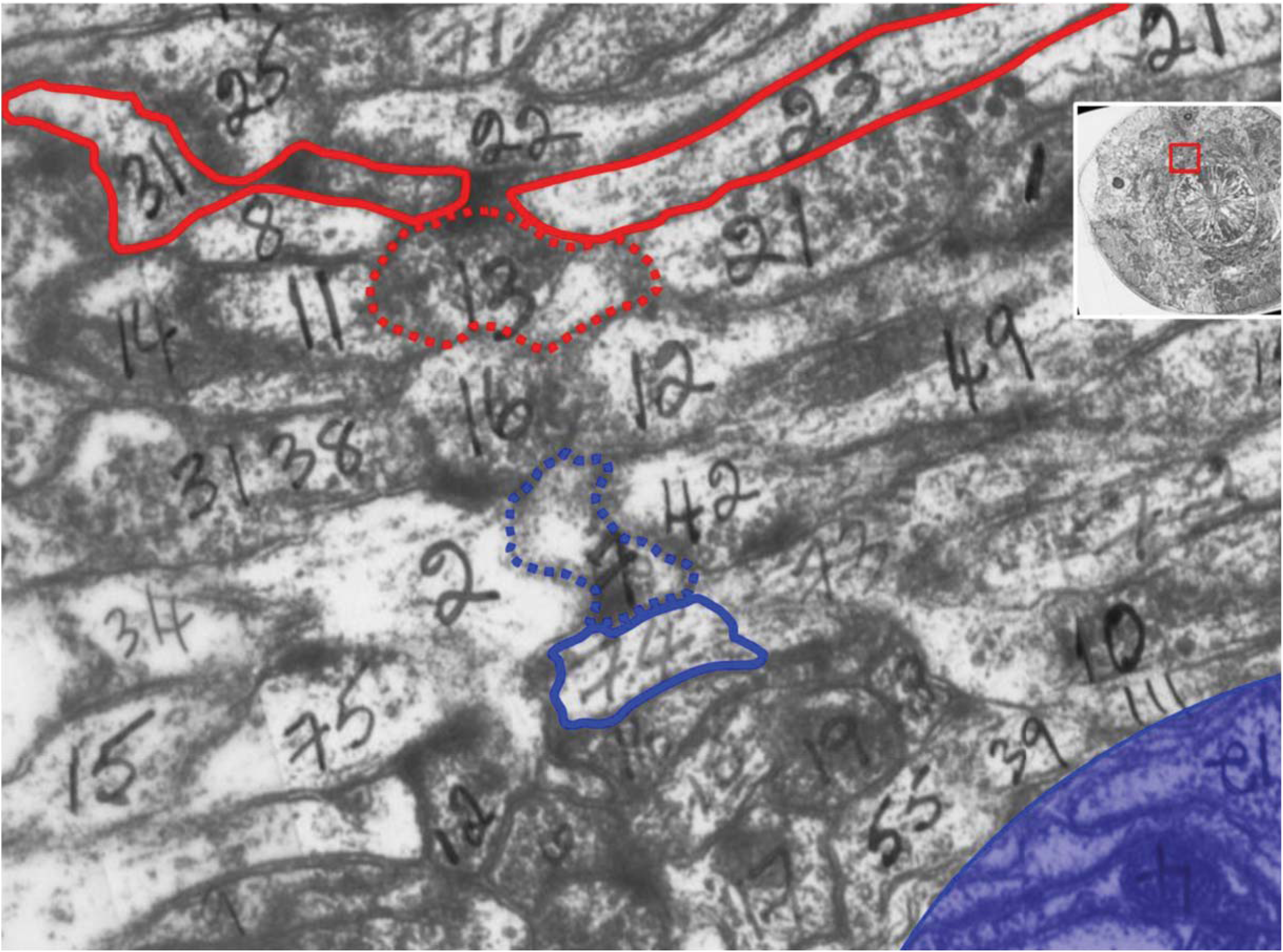
Analysis of EM data from the *C. elegans* nerve ring suggests that synapses of different circuits are found in great proximity. This figure is based on a section of the original EM image from White et al’s “The Mind of the Worm” (through WormAtlas.org and WormWiring.org). The image corresponds to slice # 87, where penciled numbers mark cell assignments. Our assignment of synapses is based on analysis by the Hall and Emmons labs, as appearing in WormWiring.org. We marked tentative cell outlines with limited accuracy (based on our best estimate from this image, and the images of adjacent slices). Cells of the avoidance circuit (ASH -> AVA/AVD/AVE) are marked with a blue outline; cells of the salt chemoattraction circuit (ASE -> AIA/AIB/AIY) are marked with a red outline. In both cases, presynaptic terminals are marked in a dashed line. The image shows a chemical synapse between presynaptic ASHL (cell #7) and postsynaptic AVDL (cell #74) and AVBL (cell #2) (although AVAL is not seen here, in the immediately following slices AVAL joins this synapse, as it squeezes between AVDL and AVBL). Another chemical synapse is formed between presynaptic ASEL (cell #13) and postsynaptic AIYL (cell #31) and AIBL (cell #23). Blue shade at the bottom right indicates the pseudocoelomic area between the nerve ring and the pharynx containing the end of the muscle arms and GLR cells. The insert in the upper right shows a zoom-out view of this area, with a red box corresponding approximately to the enlarged area. Key to cell numbers is based on cytoshow.org. Note: these numbers are only part of the full cell designation, so some numbers appear more than once. Key: 1 – AIAL; 2 - AVBL; 3 (should have been 18) - URADL; 4 (written upside down on lower right) DLV4 DBW muscle arm; 6 - URBL; 7 - ASHL; 8 – AIZL; 9 -; 10 -; 11 - ADFL; 12 (between red and blue marked synapses) – AWAL; 12 (lower center part of image) – ADAL; 12 (written upside down, lower right) - DLV12 DBW muscle arm; 13 - ASEL; 14 - RIR; 15 (lower left) – RID; 15 - OLQDL; 16 (center, between red and blue marked synapses) - AWBL; 16 (lower center) – URYDL; 17 – IL1DL; 19 – IL2DL; 21 - ASKL; 22 - AWCL; 23 - AIBL; 25 - AFDL; 31 (top left, outlined in red) – AIYL; 31 (center left, no outline) – PVPL; 34 - PVCR; 38 - AVHR; 39 - PVNL; 42 – HSNL; 49 – HSNR; 55 - PVR; 71 - AINL; 72 – AVHL; 73 AVJL; 74 - AVDL; 75 – AVJR; 111 – PVNR

To expand our study beyond slice #87 (Figure 6), we scanned further through a considerable number of slices where these synapses are found (in the range of slices #~80-130), using WormWiring.org and CytoShow.org. Supplementary figure 3 presents further annotation of slices # 79, 84, 94, and 99 as an example. We find that the ASE -> AIA/AIB/AIY synapses are consistently found closer to (and seem to have synaptic vesicles released toward) the outer rim of the nerve ring (closer to the muscle and surrounding hypodermis). In contrast, the ASH -> AVA/AVD/AVE synapses are found closer to (and seem to release synaptic vesicles toward) the inner rim of the nerve ring (closer to the pharynx). In our previous studies (Mano et al., 2007), the expression of *glt-4* was assigned to neurons, but neuronal identities were difficult to ascertain, obscuring the basis for its privileged role. In contrast, both partial (Mano et al., 2007) and full-length protein fusion (data not shown) of GLT-1::GFP indicate that *glt-1* is heavily expressed in head muscles and in hypodermis, while Katz & Shaham show that it is also expressed in cephalic sheath glia (Katz et al., 2018) that wraps around the outer circumference of the nerve ring. The juxtaposition of the ASE -> AIA/AIB/AIY synapses and the *glt-1* (proximal GluT) -expressing head muscles, hypodermis, and glia could therefore provide a basis for the privileged role of *glt-1* in clearing Glu released from ASE, and explain the susceptibility of *glt-1* KO animals to the putative spillover of Glu out of ASE -> AIA/AIB/AIY synapses (as evidenced by AVA stimulation by low NaCl concentrations in these animals, Figures 4 & 5).

### Motility of the head and pharynx is critical to preserve the fidelity of AVA synaptic activity

Although the privileged role of the proximal *glt-1* on ASE -> AIA/AIB/AIY synapses can now be reasonably explained by the proximity of these synapses to the head muscles, hypodermis, and glia, it remains unclear how the distal GluTs (*glt-3, glt-6*, & *glt-7*), expressed on the canal cell, might preferentially affect the ASH -> AVA/AVD/AVE synapses, which are closer to the pharynx. Since the canal cell is directly exposed to pseudocoelomic body fluids, distal GluTs are likely regulate ambient Glu levels in body fluids. Ambient extracellular Glu concentrations in the vicinity of mammalian synapses are known to affect synaptic Glu clearance by diffusion (Kullmann and Asztely, 1998; Bergles et al., 1999; Diamond, 2002). Interestingly, in the worm, nerve ring interstitial fluids are continuous with the pseudocoelomic fluid found in the space between the nerve ring and the isthmus of the pharynx (marked as blue shaded arch in Figure 6 and in Supplementary figure 3). This fluid compartment is connected to the rest of the pseudocoelomic body fluids (see https://www.wormatlas.org/hermaphrodite/introduction/mainframe.htm#IntroFIG2 and http://wormatlas.org/hermaphrodite/pericellular/Periframeset.html). According to WormAtlas, the pseudocoelomic fluid in this region is believed to reduce friction between adjacent tissues arising from vigorous pharyngeal movements during feeding. Solutes in fluid of this region seem to have unfettered access to the inner rim of the nerve ring; the mesh-like basal laminae that separate different tissues are fully permeable to neurotransmitters and other small molecules (Kandel et al., 2013). We therefore considered a possible link between the putative ability of distal GluTs to control ambient Glu concentration in pseudocoelomic body fluids and the location of AVA synapses closer to the inner rim of the nerve ring. Considering that our experiments thus far were performed on physically-restrained (but not pharmacologically-paralyzed) animals, we contemplated the possibility that the continuous pulsations of the pharynx and the increased mobility that nematodes show in the head region (causing the nerve ring to slide slightly back and forth on the isthmus, even in restrained animals), may facilitate perfusion of body fluids to the inner part of the nerve ring. Such perfusion can replace extracellular fluids rich in synaptically released Glu with fresh body fluids containing lower, ambient levels of Glu. If normal Glu clearance from the inner rim is aided by mechanical agitation and perfusion, then paralysis might increase ambient Glu levels, potentially hampering Glu diffusion from these synapses, especially under conditions of increased synaptic activity and reduced Glu clearance. We therefore hypothesize that mechanical agitation of body fluids and perfusion of the inner rim of the nerve ring might underlie the privileged effect of distal GluTs on the ASH -> AVA synapses (an effect seen in Figures 1-3) and affect the clearance of Glu that reach AVA by spillover (Figures 4 and 5). In line with this hypothesis, we recently saw that under conditions where Glu accumulation leads to nematode excitotoxicity (Mano and Driscoll, 2009), the neurons that are most severely affected by neurodegeneration are not those that express the most GluRs, but those who form Glu synapses in the innermost face of the nerve ring (Feldmann et al., 2019).

Further to this hypothesis, we suspect that inhibiting pharyngeal pumping and animal motility may prevent access of fresh body fluids to the synapses (in both WT and GluT KO animals), and interfere with the activity of synapses that depend on it. To test this hypothesis we first used tetramisole (Lewis et al., 1980; Lewis et al., 1987), which works as a constitutive, desensitizing agonist of nicotinic AcetylCholine Receptors (nAChRs) and is routinely used to paralyze worms for GCaMP-based imaging of specific neurons (Hendricks et al., 2012; Larsch et al., 2013; Kato et al., 2014) or the whole nervous system (Schrödel et al., 2013). However, we had some reservations on using tetramisole in our specific studies because of nAChR expression in AVA (Feng et al., 2006; Sherlekar et al., 2013), though spontaneous activity and indirect odor responses of AVA remain intact in the presence of tetramisole (Schrödel et al., 2013; Gordus et al., 2015). To demonstrate the effect of paralysis without disrupting neuronal activity, we also performed experiments using BDM (Goodman and Chalfie, 1998), which induces paralysis directly in the muscle by inhibiting myosin. Though BDM also has off-target effects, we speculate that similar effects seen when using either tetramisole or BDM are most likely to come from their common effect on paralysis.

Prior to our paralysis experiments, we needed to run a few controls. We first verified that the paralyzing agent, and especially the neuronally-active tetramisole, does not affect the activity of the presynaptic neuron. Indeed, by using GCaMP expressed in ASH we verified that tetramisole did not diminish ASH activity (Supplementary Figure 4). Secondly, we verified that a paired stimulation separated by 10 minute recovery shows no diminution of AVA response to the second stimulus. We performed this analysis for both AVA response to ASH stimulation with glycerol and for ASE stimulation with low concentration NaCl, confirming that the previously described AVA responses in the different GluT KOs (Figures 2 and 5) are preserved in paired stimuli of non-paralyzed animals (Supplementary figure 5).

To test the effect of paralysis on AVA responses to either ASH or ASE stimulation, we first applied stimulus under normal conditions, establishing the normal response in AVA in the specific animal. We then paralyzed the worm anterior with either tetramisole or BDM in absence of chemical stimulation for 10 minutes. Finally, we exposed the paralyzed animal to the same stimulus a second time. We found that paralysis with tetramisole eliminated the exaggerated response of AVA to ASH stimulation in *glt-3, glt-6, glt-7* KO mutants (Figure 7). Similarly, tetramisole-mediated paralysis abolished the response of AVA to ASE stimulation in *glt-1* and *glt-4* mutants (Figure 8). We observed similar effects when worms were paralyzed with BDM (Supplementary Figures 6 and 7), suggesting the effects we see do not arise from the various side effects of the two drugs, but from the shared effect of paralysis. If paralysis halts perfusion-mediated Glu clearance around AVA, then normal spontaneous activity (without stimulation) may result in Glu accumulation in AVA synapses during prolonged incubation with the paralyzing agent. In support of this hypothesis, we find GCaMP fluorescence in AVA to increase shortly after the onset of paralysis (Supplementary figure 8). However, we could not resolve the details of this effect with direct Glu measurements at this time, because the weak iGluSnFR signals in AVA neurites are not conducive to such protracted imaging.

**Figure 7:**
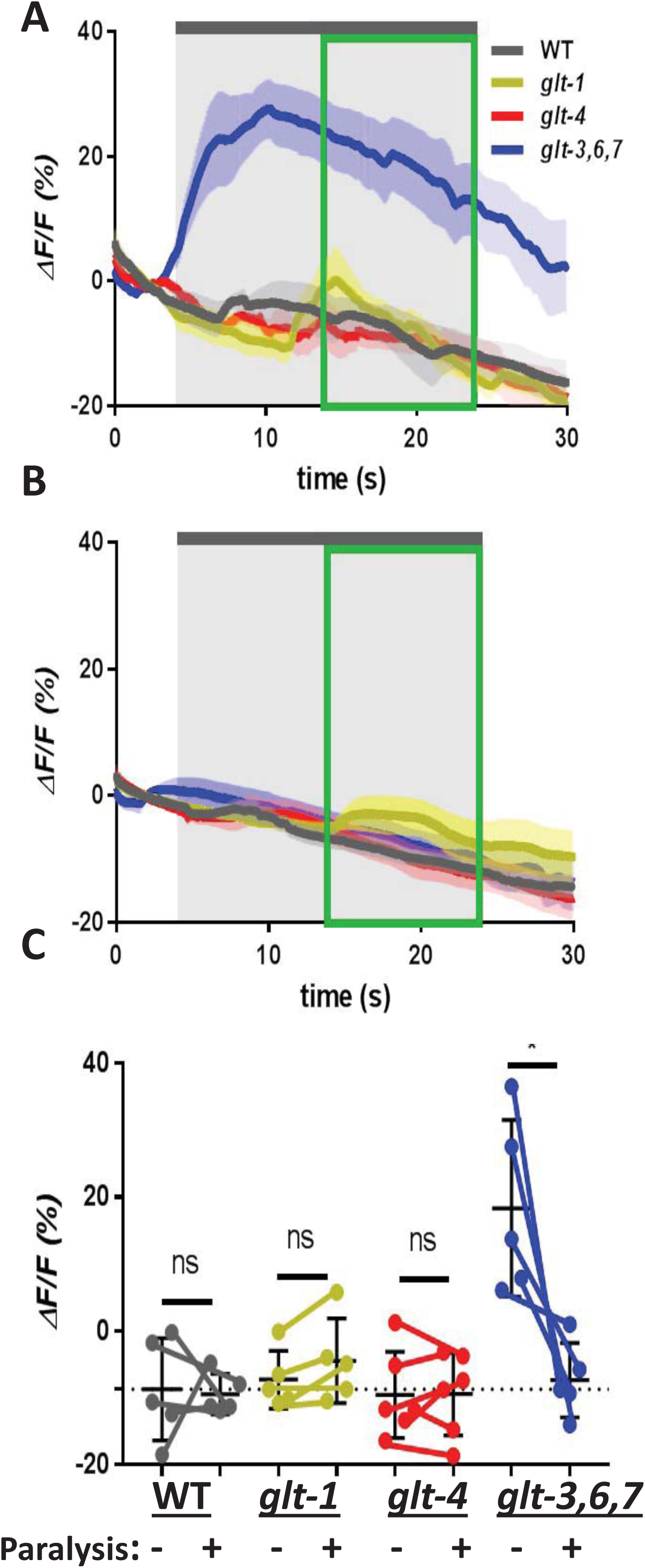
Tetramisole-induced paralysis eliminates exaggerated responses in the ASH -> AVA synapse to 1 M glycerol. Head muscle and pharyngeal paralysis induced by exposure to tetramisole caused loss in AVA response of *glt-3,6,7* mutants to stimulation by 1 M glycerol. Changes in AVA GCaMP fluorescence intensity in response to 1 M glycerol stimulation are shown before **(A)** and following **(B)** 10 min paralysis treatment with tetramisole. Traces are labeled as in previous slides, but steady state response of each animal was calculated separately. **(C)** Paired comparison of steady-state responses in individual animals before and following paralysis are shown as line-connected dots. The average of responses in each group appears as a horizontal thin bar with error bars. Dotted horizontal line represents the average response of WT animals before paralysis, to which the other responses are compared. Statistical significance is denoted with asterisks. * *P* = 0.0283, ANOVA with Bonferroni correction. n = 5-6 per strain.

**Figure 8:**
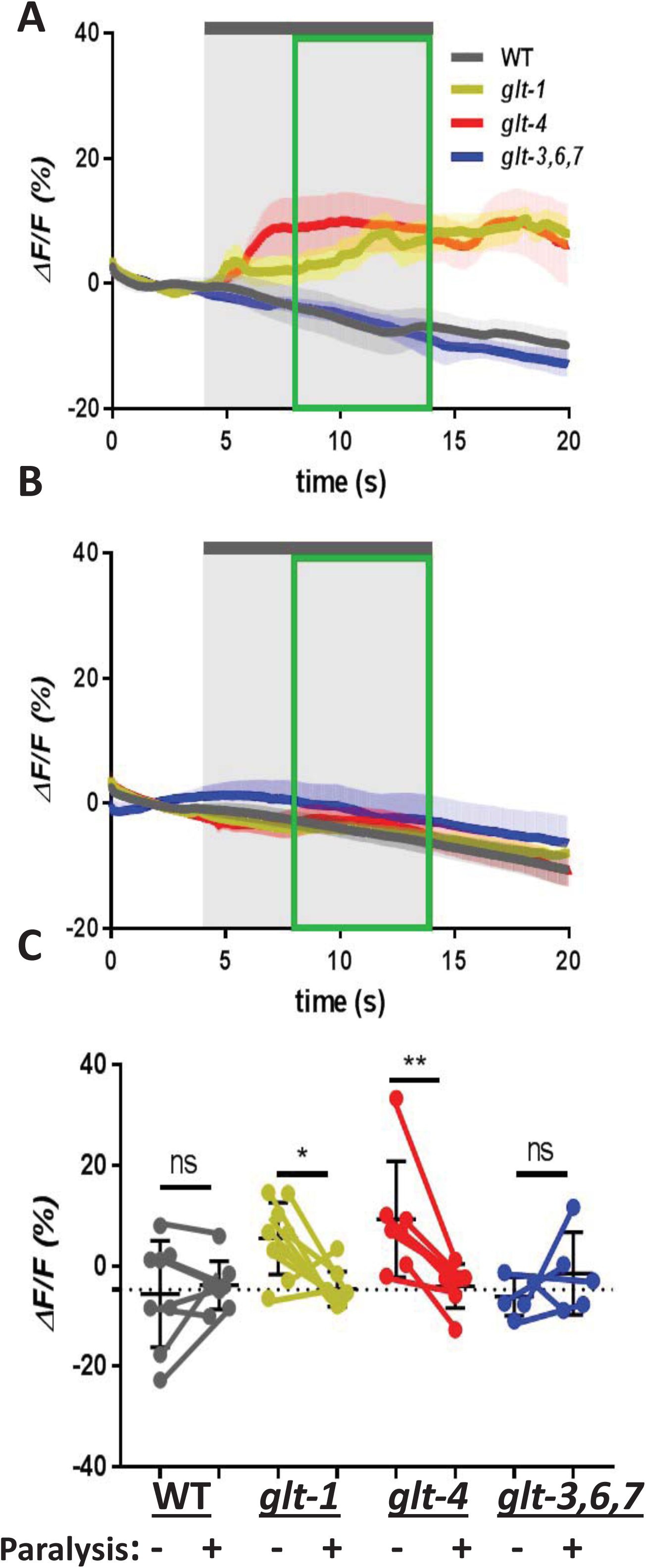
Putative spillover from ASE onto AVA is eliminated by tetramisole-induced head paralysis. Head muscle and pharyngeal paralysis induced by exposure to tetramisole caused loss in AVA response of *glt-1* and *glt-4* mutants to stimulation by 1 mM NaCl. Changes in AVA GCaMP intensity in response to 1 mM NaCl stimulation analyzed as in figure 7. * *P* = 0.0132; ** *P* = 0.0077, ANOVA with Bonferroni correction. n = 5-9 per strain.

In summary, our observations establish that the ability of the worm to continuously agitate interstitial / pseudocoelomic fluids is a critical factor in normal physiology of Glu signaling in some synapses of the aspiny and glia-deprived synaptic hub of the nematode. Additionally, this explains how distal GluTs display a privileged role in clearance of synapses located near the inner rim of the nerve ring.

## Discussion

Our study focuses on synaptic clearance of Glu, a widely used non-degradable neurotransmitter, in a compact nervous system lacking spines and glia-mediated anatomical separation between synapses. We find that in the nematode nervous system, synaptic fidelity and circuit resolution is maintained by a robust, two-tier system of proximal and distal Glu clearance. We observe that specific GluTs have a privileged role in preserving accuracy and preventing spillover in specific circuits. We affirm that the privileged role of distal GluTs (*glt-3, glt-6*, & *glt-7*) on the ASH -> AVA/AVD/AVE avoidance circuit, seen previously at the behavioral level (Mano et al., 2007), can be observed at the neurophysiological level as hyperactivation of somatic Ca^2+^ responses and dendritic Glu increase in postsynaptic AVA neurons (Figures 1, 2, and 3). Distal GluTs expressed on canal cells, through direct contact with pseudocoelomic body fluids, putatively maintain low ambient Glu concentrations to enable effective diffusion-mediated clearance. Although initially it was unclear to us how body fluids access these synapses, the notion of diffusion-mediated distal clearance in nematodes is in line with the anatomical observation that nematode synapses are aspiny and formed *en passant* (White et al., 1986), promoting perisynaptic diffusion. In contrast to the functional association of ASH synapses with distal GluTs, the accuracy of ASE -> AIA/AIB/AIY synapses in the low-salt sensing chemoattraction circuit is more influenced by the activity of the proximal GluTs, encoded by *glt-1* and *glt-4* (Figures 3 & 5): in the absence of these proximal GluTs, ASE circuit resolution is lost and Glu seems to spillover onto nearby circuits. The localization of ASE -> AIA/AIB/AIY synapses to the outer rim of the nerve ring (Figure 6 and Supplementary figure 3) offers a reasonably straightforward explanation to the privileged role of proximal GluTs on these synapses, as these synapses are closer to the *glt-1* – expressing hypodermis, head muscles, and glia. Spillover from other circuits into ASE synapses has yet to be examined, and remains an open question.

The location of ASH -> AVA/AVD/AVE synapses closer to the inner rim of the nerve ring and their functional association with distal GluTs (maintaining ambient Glu concentrations) is particularly intriguing. Careful examination of anatomical data in WormAtlas brought an important notion to our attention, namely that the space between the inner rim of the nerve ring and the isthmus of the pharynx is filled with pseudocoelomic body fluids, a compartment linked to fluids in the rest if the body. We propose that together with the intense mechanical agitation in this area, these body fluids might provide perfusion of the inner rim of the nerve ring, which is contiguous with the interstitial fluids in the neuropil extracellular space. Indeed, we observe that inhibiting fluid agitation (using two different paralyzing agents) obstructs chemical stimulation of AVA (either directly from ASH or by spillover from ASE, Figures 7 & 8). This notion is further supported by our recent observation that under conditions that cause *glt-3* KO–induced excitotoxicity, the most severely affected neurons are not those that express the most GluRs, but rather those that face the innermost rim of the nerve ring (Feldmann et al., 2019).

In our model for the two-tier design of the Glu clearance system in *C. elegans*, Glu secreted closer to the outer rim of the nerve ring is preferentially cleared by GluTs expressed on the large structures that surround the nerve ring (hypodermis, head muscles, and glia), while Glu released closer to the inner rim of the nerve ring is preferentially cleared by circulating body fluids and distal uptake into the canal cell by distal GluTs (Supplementary figure 9). In this study, we did not investigate where Glu synapses of other circuits are located and how they are cleared, nor do we know if this two-tier organization of synaptic clearance is unique to Glu or is shared by other neurotransmitters. However, a broader applicability of this clearance mechanism is possible, since a number of neurotransmitters in *C. elegans* have been suggested to spill from their synapse of origin, and are therefore also candidates for long-range clearance (Chase et al., 2004; Jafari et al., 2011; Jobson et al., 2015). There is considerable likelihood for this mechanism to affect additional neurotransmitters since the canal cell expresses additional neurotransmitter transporters, such as the betaine/GABA transporter *snf-3* (Peden et al., 2013).

The incomplete isolation of synapses throughout animal phyla suggests that in other animals, circuit resolution may be maintained not only by anatomical separation, but also by a balance between physical isolation and functional means of Glu clearance from both synaptic and perisynaptic or interstitial spaces. Our results on the physiology of Glu signaling and clearance in the nematode suggest this balance between anatomical and physiological means effectively prevents spillover and ensures signaling accuracy. In this view, vigorous clearance of Glu can compensate for anatomical shortcoming caused by the lack of glia isolation of synapses, as the existence of this two-tier Glu clearance system allows for preservation of circuit resolution in the nematode, even in the absence of anatomical synaptic isolation. Most intriguingly, we present data to support a hypothesis that agitation of interstitial fluids and mechanical perfusion might be a considerable factor in Glu clearance in some key synapses in the nematode, allowing for subsequent clearance by distal GluTs.

It is interesting to note the correlation between synaptic structure and its physiology. Flat synapses such as those seen in the nematode’s aspiny neurons are believed to be particularly efficient in passive cable propagation of receptor potential (Segal, 2010). This may hold special significance in the context of the (mostly) non-spiking graded signaling seen in the nematode’s nervous system and the lack of a Mg^2+^ block in its NMDA-Rs (Brockie et al., 2001b), eliminating the “need” for dendritic spines in this animal. These structural characteristics seem to align with a clearance strategy that relies more heavily on clearance by diffusion and perfusion. It will therefore be interesting to study the effect of perfusion in the nervous systems of higher animals, which possess a complete spectrum of synaptic morphologies, glial involvement (Harris, 1999; Segal, 2010; Thomas et al., 2011), and a range of levels of exposure to the glymphatic system (Nicholson and Hrabetova, 2017; Da Mesquita et al., 2018; Rasmussen et al., 2018).

It is also worth noting that pulsations of mammalian brain parenchyma and CSF is readily observed (Hadaczek et al., 2006; Wagshul et al., 2011). In the murine brain, pulsatility of cerebral arteries has been recently shown to enhance interstitial fluid perfusion and is suggested to augment clearance of extracellular solutes (Iliff et al., 2013; Mestre et al., 2018). Pulsatility has been also used to preserve brain function after damage (Cohn et al., 2015; Vrselja et al., 2019). Furthermore, reduction of Glu clearance by agitation of interstitial fluid might have a pronounced role in brain damage during pulsation disturbances, and in conditions where extracellular space in the neuropil is reduced, such as brain edema (Sherpa et al., 2014) and other pathological conditions (Arbel-Ornath et al., 2013). Such disruptions might selectively affect glia-deprived flat synapses, where clearance by diffusion and bulk flow may be more critical. Continued study of the functional organization of neurotransmitter clearance in the nematode nervous system has the potential to elucidate unexpected and broadly applicable basic principles in the physiology of synaptic clearance. These insights might broaden the discussion of mechanisms that maintain accuracy of synaptic signaling and the resolution of neuronal circuits.

## Authors contribution

IM, JC & KKL, designed the project, analyzed the data, and wrote the manuscript, JC & KKL performed all the imaging experiments, JCYW and PM performed the behavioral experiments.

