## Supplementary material for "Novel Strategies for Glutamate Clearance in the Glia-Deprived Synaptic Hub of *C. elegans*": Supp

*Abbreviated Title: Glutamate Clearance in Nematode Glia-Deprived Synapses*

Joyce Chan, Kirsten KyungHwa Lee, Jenny Chan Ying Wong, Paola Morocho, and Itzhak Mano.

**Supplementary figure 1**

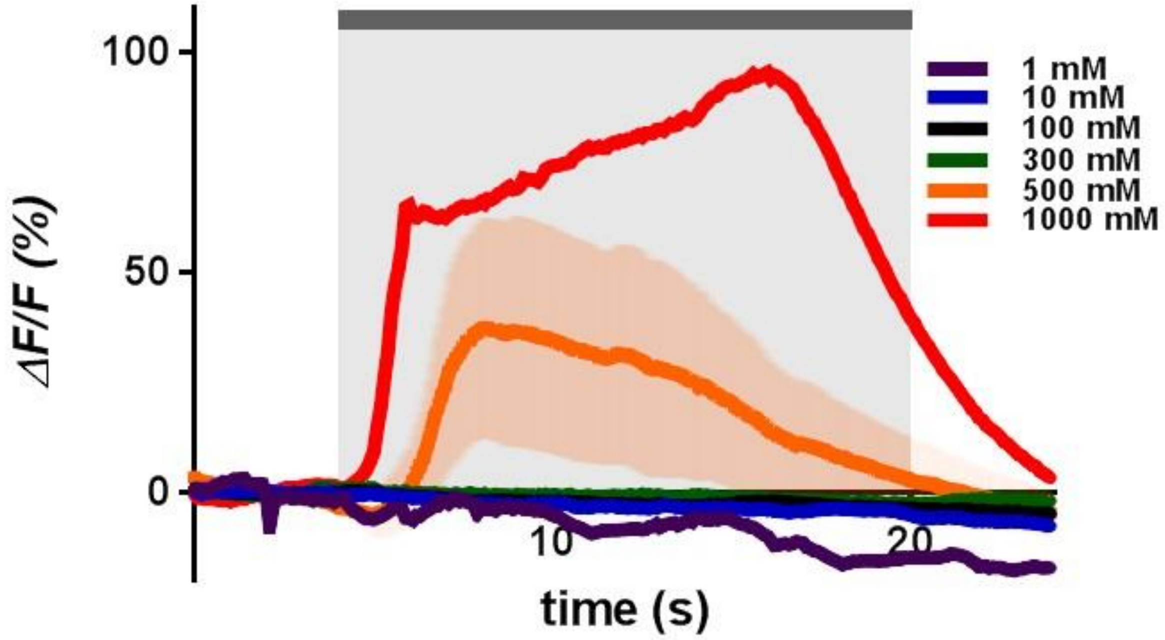


**ASH neurons respond only to high concentration NaCl.**

NaCl-induced calcium responses in ASH neurons of WT animals, showing traces of average changes in GCaMP3 fluorescence. Shaded areas above and below the trace represent SEM. The fluorescence change for the first 4 s before stimulation was averaged to serve as the baseline fluorescence, $F_0$. Light gray shading indicates 10 s of NaCl application at each concentration, beginning at *t* = 4 s. n = 2-3 per concentration.

**Supplementary figure 2**

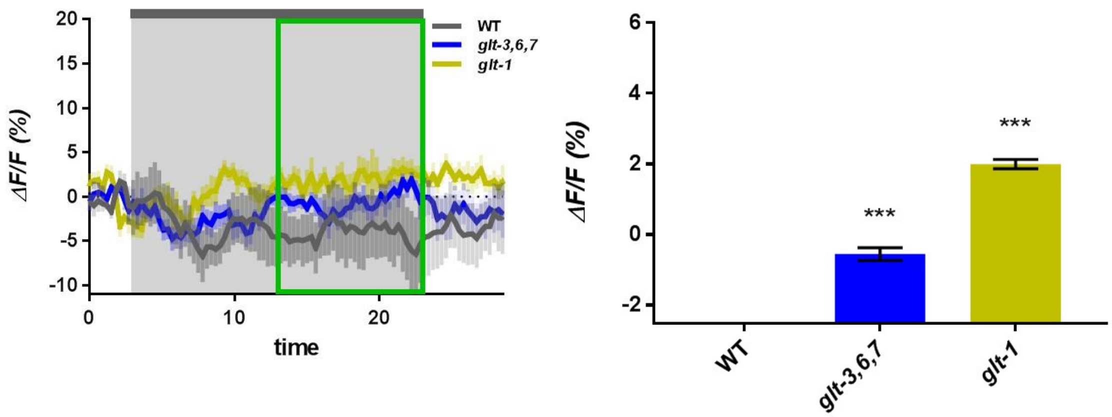


**Proximal GluT KOs promote accumulation of Glu at the AVA synapse following 1mM NaCl stimulation.**

NaCl-induced changes in Glu concentrations in nerve ring processes of AVA neurons in WT and GluT KO animals. Average traces of changes in iGluSnFR fluorescence and averaged steady-state responses to a 1mM NaCl stimulus are analyzed similarly to those in Figure 2. *** $P < 0.001$, ANOVA with Bonferroni correction. n = 5-7 for each strain.

**Supplementary figure 3**

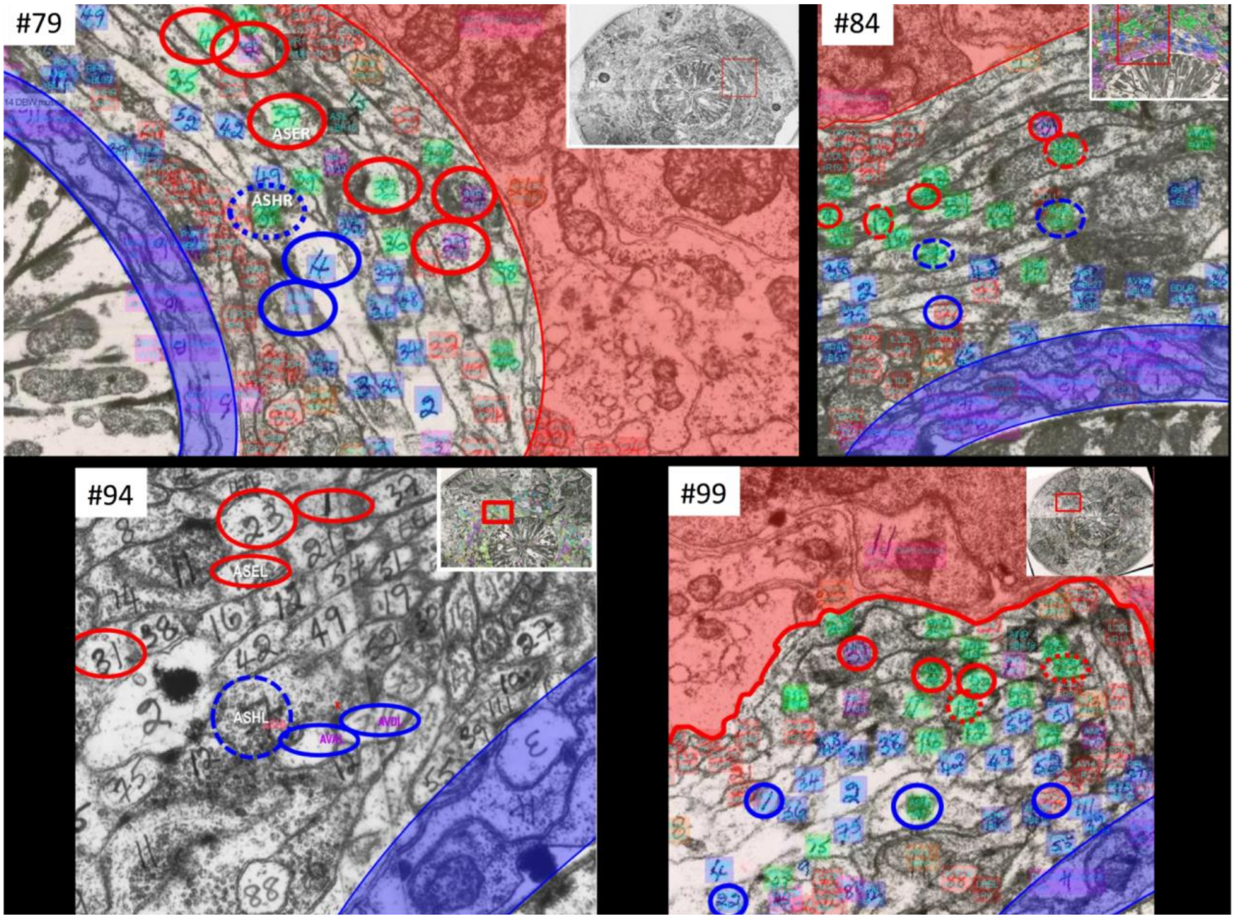


**Analysis of EM data from the *C. elegans* nerve ring suggests that synapses of different circuits are found in great proximity.**

The figure is based on a section of the original EM image from White *et al*'s “The Mind of the Worm” [White, 1986 #1816] (through WormAtlas.org and WormWiring.org). The image corresponds to slice # 87, where penciled numbers mark cell assignments. Our assignment of synapses is based on analysis by the Hall and Emmons labs, as appearing in WormWiring.org. We marked tentative cell outlines with limited accuracy (based on our best estimate from this image, and images of adjacent slices). Cells of the avoidance circuit (ASH -> AVA/AVD/AVE) are marked with a blue outline; cells of the salt chemoattraction circuit (ASE -> AIA/AIB/AIY) are marked with a red outline. In both cases, presynaptic terminals are marked in a dashed line. The image shows a chemical synapse between presynaptic ASHL (cell #7) and postsynaptic AVDL (cell #74) and AVBL (cell #2) (although AVAL is not seen here, in the immediately following slices AVAL joins this synapse, as it squeezes between AVDL and AVBL). Another chemical synapse is formed between presynaptic ASEL (cell #13) and postsynaptic AIYL (cell #31) and AIBL (cell #23). Blue shading at the bottom right indicates the pseudocoelomic area between the nerve ring and the pharynx containing the end of the muscle arms and GLR cells. The insert in the upper right shows a zoom-out view of this area, with a red box corresponding approximately to the enlarged area.

Key to cell numbers is based on cytoshow.org . Note: these numbers are only part of the full cell designation, so some numbers appear more than once. Key: 1 – AIAL ; 2 - AVBL ; 3 (should have been 18) - URADL ; 4 (written upside down on lower right) DLV4 DBW muscle arm ; 6 - URBL ; 7 - ASHL ; 8 – AIZL ; 9 - ; 10 - ; 11 - ADFL ; 12 (between red and blue marked synapses) – AWAL ; 12 (lower center part of image) – ADAL ; 12 (written upside down, lower right) - DLV12 DBW muscle arm ; 13 - ASEL ; 14 - RIR ; 15 (lower left) – RID ; 15 - OLQDL ; 16 (center, between red and blue marked synapses) - AWBL ; 16 (lower center) – URYDL ; 17 – IL1DL ; 19 – IL2DL ; 21 - ASKL ; 22 - AWCL ; 23 - AIBL ; 25 - AFDL ; 31 (top left, outlined in red) – AIYL ; 31 (center left, no outline) – PVPL ; 34 - PVCR ; 38 - AVHR ; 39 - PVNL ; 42 – HSNL ; 49 – HSNR ; 55 - PVR ; 71 - AINL ; 72 – AVHL ; 73 - AVJL ; 74 - AVDL ; 75 – AVJR ; 111 – PVNR.

**Supplementary figure 4**

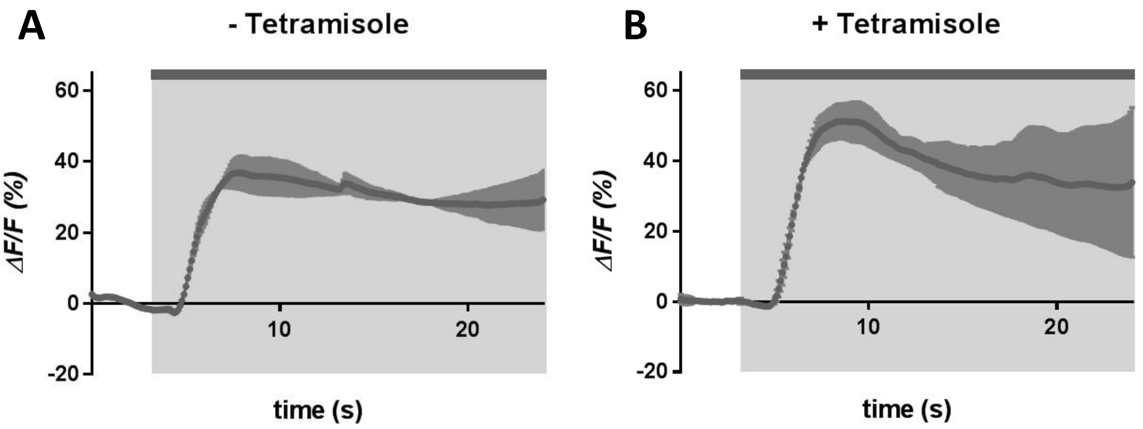


**ASH responds to 1 M Glycerol before and during paralysis with 2 mM tetramisole.** WT animals expressing GCaMP in AVA neurons were imaged before (**A**) and during paralysis (**B**) induced by 2 mM tetramisole**.** Light gray rectangle in **A** and **B** indicates the 20 s period of stimulation with 1 M glycerol, beginning at *t* = 4 s. Dark gray shading around each curve indicate SEM. n = 2.

**Supplementary figure 5**

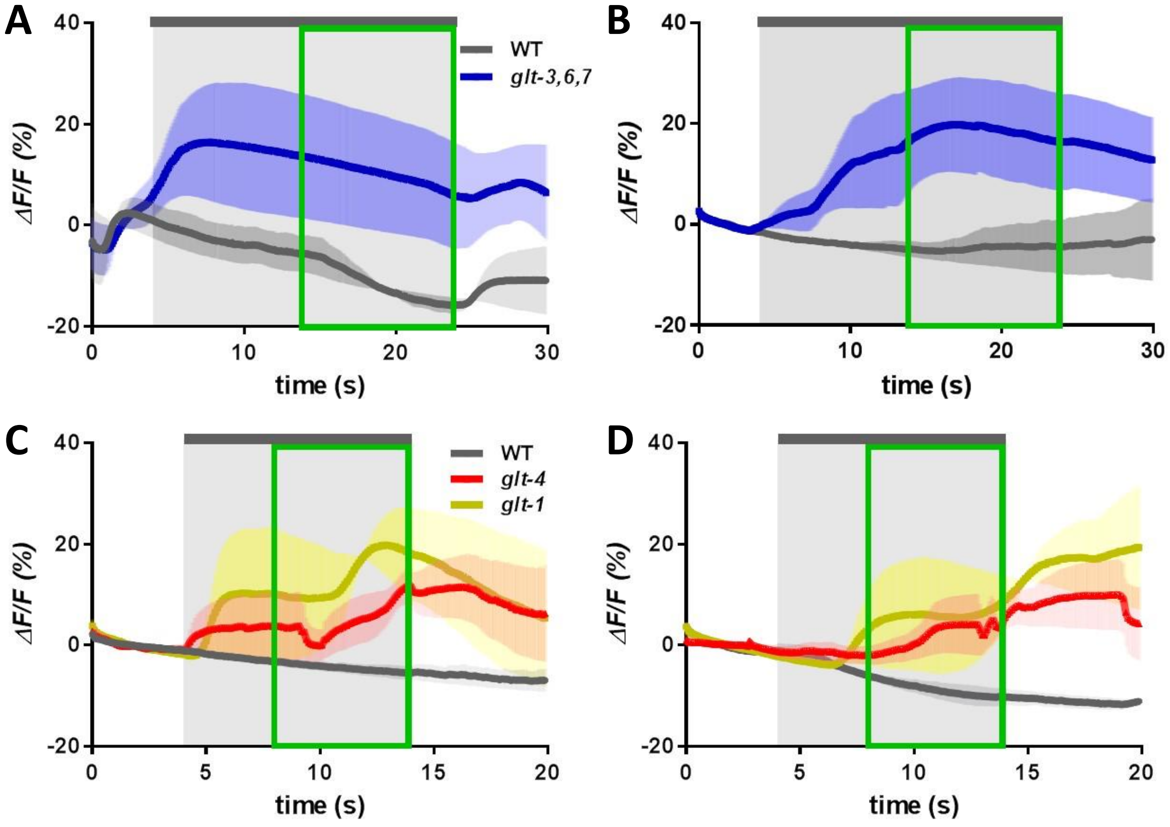


**Sequential stimulation of worms with 1 M glycerol or 1 mM NaCl does not diminish the exaggerated responses in AVA or putative spillover from ASE**.

Calcium responses in AVA neurons of WT and GluT KO animals before **(A, C)** and following **(B, D)** 10 min sham incubation with buffer. Average traces of changes in GCaMP3 fluorescence and averaged steady-state responses to either **(A, B)** 1 M glycerol or **(C, D)** 1 mM NaCl stimuli are analyzed similarly to those in Figure 2. n = 2-4 for each strain.

**Supplementary figure 6**

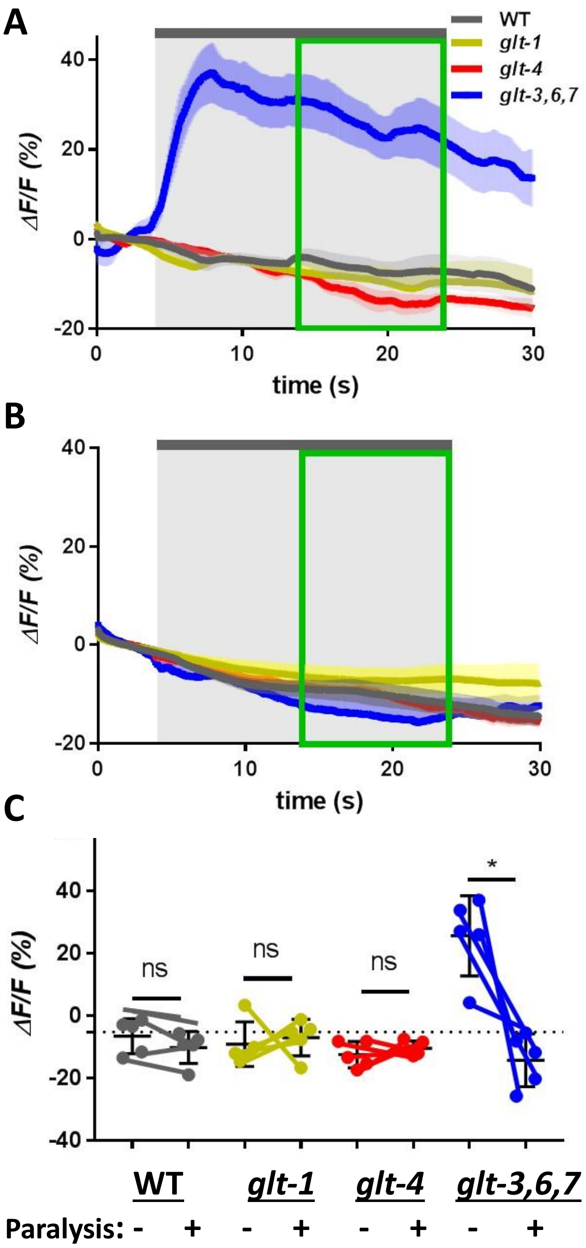


**BDM-induced paralysis eliminates exaggerated responses in the ASH -> AVA synapse to 1 M glycerol.** Head muscle and pharyngeal paralysis induced by exposure to BDM caused loss in AVA response of *glt-3,6,7* mutants to stimulation by 1 M glycerol. Changes in AVA GCaMP fluorescence intensity in response to 1 M glycerol stimulation are shown before **(A)** and following **(B)** 10 min paralysis treatment with BDM. Traces are labeled as in previous slides, but with separate steady state response calculations for each animal. **(C)** Paired comparison of steady-state responses in individual animals before and following paralysis are shown as data points connected by lines. The average of responses in each group appears as a horizontal thin bar flanked by SEM error bars**.** Dotted horizontal line represents the average response of WT animals before paralysis, to which the other responses are compared. Statistical significance is denoted with asterisks. * *P* = 0.0106, ANOVA with Bonferroni correction. n = 5 per strain.

**Supplementary figure 7**

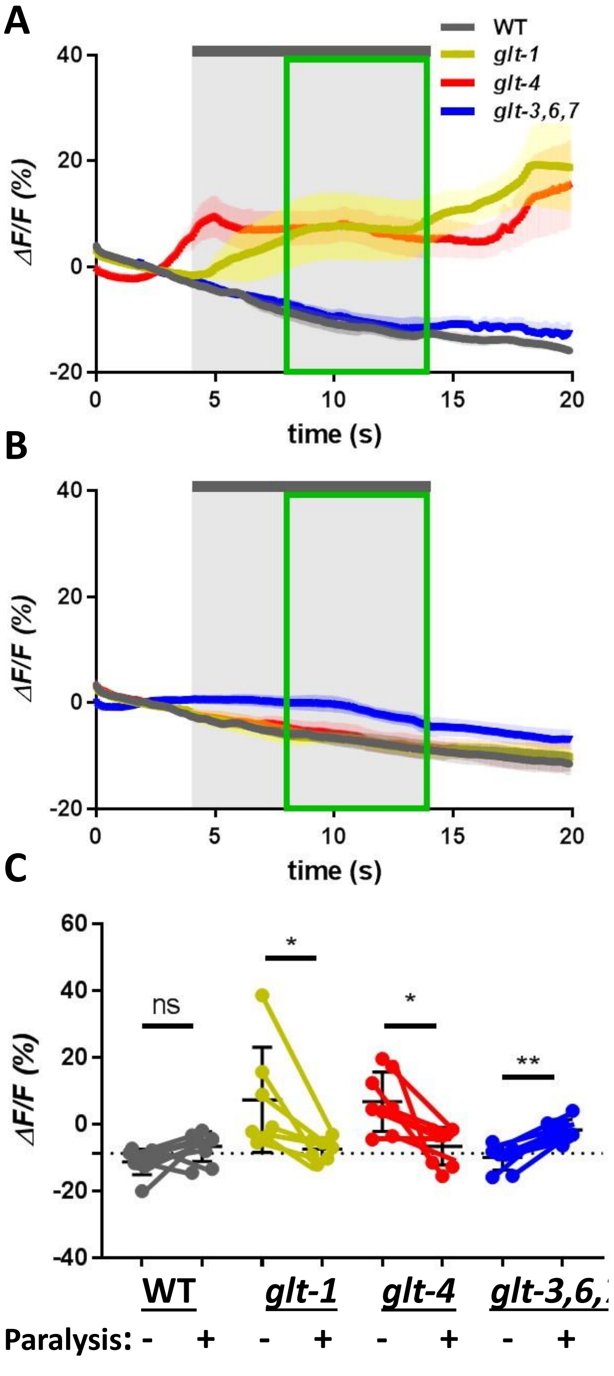


**Putative spillover from ASE onto AVA is eliminated by BDM-induced head paralysis.**

Head muscle and pharyngeal paralysis induced by exposure to BDM caused loss in AVA response of *glt-1* and *glt-4* mutants to stimulation by 1 mM NaCl. Changes in AVA GCaMP intensity in response to 1 mM NaCl stimulation analyzed as in supplementary figure 6. * *P* = 0.0330 for *glt-1* and 0.0145 for *glt-4* ; ** *P* = 0.0016, ANOVA with Bonferroni correction. n = 7-9 per strain.

**Supplementary figure 8**

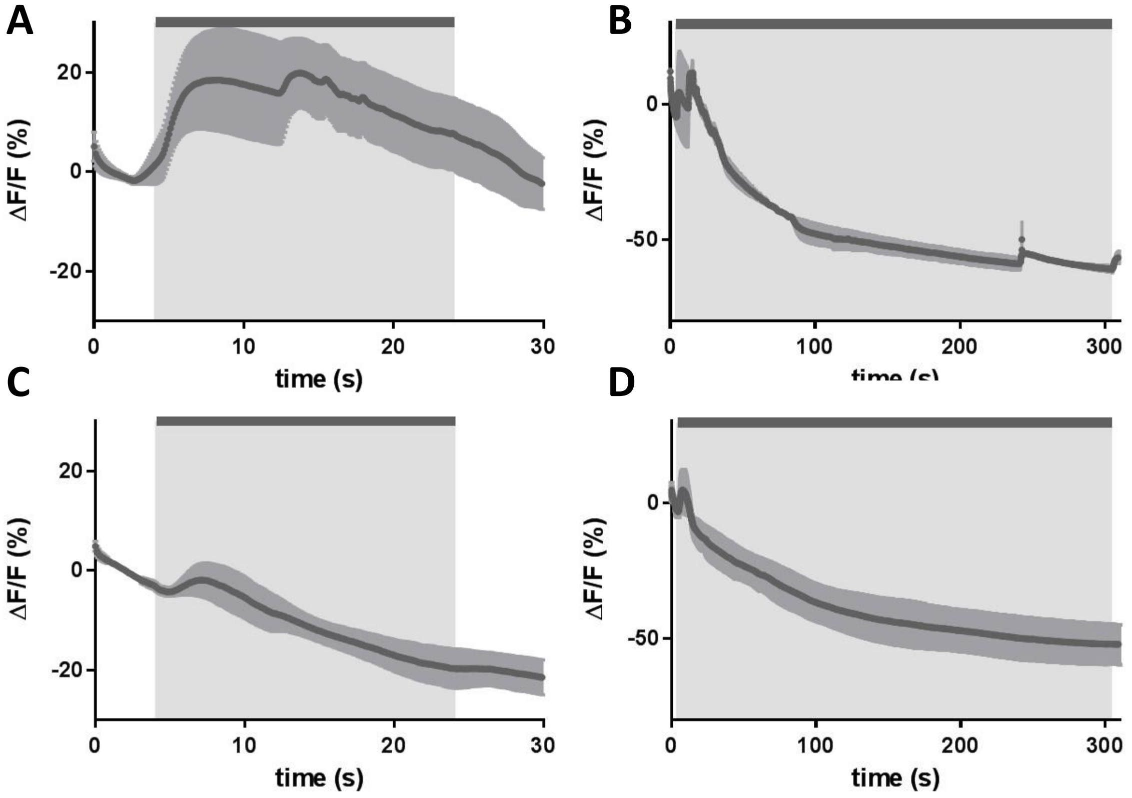


**AVA displays response upon initial exposure to 2 mM tetramisole.**

Traces of average calcium fluorescence in WT animals expressing GCaMP in AVA neurons, imaged during short **(A, C)** or long **(B, D)** exposure to tetramisole **(A, B)** or BDM **(C, D)** without stimulation. Light gray shaded box in **A** and **C** indicates initial 20 s period of exposure to the paralyzing drug; **B** and **D** display a 5 min prolonged exposure to the paralyzing drug. Shaded areas above and below the trace indicate SEM. n = 2-5.

**Supplementary figure 9**

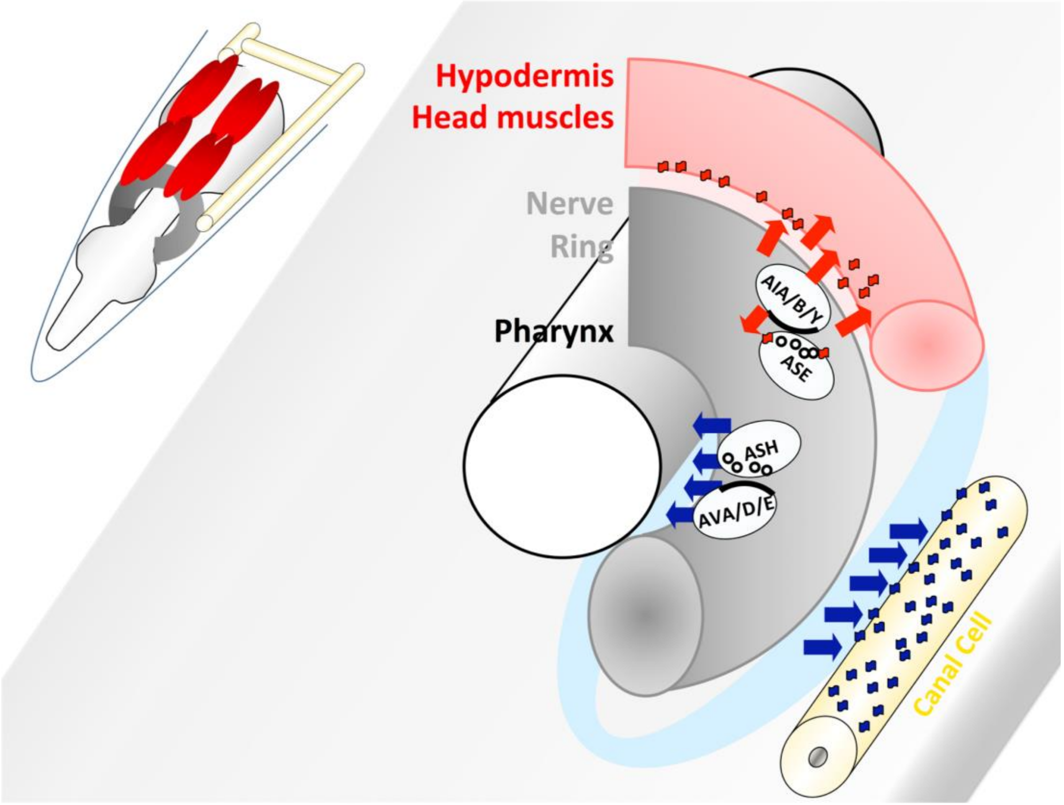


**The two-tier model of Glu clearance in *C. elegans.***

Glu released in synapses that are closer to the outer rim of the nerve ring (such as in the ASE-> AIA/AIB/AIY circuit) is preferentially cleared by proximal GluTs expressed in head muscles (red), hypodermis, and glia (not shown). Glu released in synapses that are closer to the inner rim of the nerve ring (such as in the ASH -> AVA/AVD/AVE circuit) is preferentially cleared by diffusion to interstitial fluids that are continuous with the pseudocoelomic area between the inner rim of the nerve ring and the pharynx. Glu concentration in the pseudocoelomic fluid is kept low by distal GluTs on the canal cell. Agitation by pharyngeal activity and head movement might help circulate the pseudocoelomic fluid and perfuse the inner face of the nerve ring with fresh pseudocoelomic fluid that contains low Glu concentrations.
